# An extended N-terminus restrains the plant cell death-inducing ability of the catalytically competent ribonuclease domain in a pea powdery mildew RALPH effector

**DOI:** 10.64898/2026.08.13.744637

**Authors:** Debashish Sahu, Puja Ghosh, Smritilekha Mukherjee, Vineet Kumar, Gunjan Sharma, Megha Gupta, Gagan Gupta, Poonam Ray, Kusum, Jaya Sharma, Deepti Jain, Divya Chandran

## Abstract

RALPH (RNase-like proteins associated with haustoria) effectors, which are preferentially expressed in haustoria and structurally resemble fungal T1/F1 RNases, constitute one of the largest effector families in powdery mildew (PM) fungi, yet their functions in dicot-adapted PM species remain poorly understood. Unlike cereal PM RALPHs, which lack the catalytic residues required for RNase activity, some dicot PM RALPHs retain these residues. Here, we performed a comprehensive structural and expression-based characterization of the pea PM *Erysiphe pisi* RALPH (*Ep*RALPH) repertoire and functionally characterized *Ep*RALPH11, a RALPH effector with partial conservation of the catalytic residues of T1/F1 fungal RNases. Comparative analyses identified multi-RNase-domain RALPHs as a conserved feature of the *Erysiphe* lineage, while expression profiling showed that many *EpRALPH*s are preferentially expressed in haustoria during early host colonization. AlphaFold 3-based structural analyses revealed a conserved T1/F1 RNase-like fold despite substantial sequence and surface charge divergence, indicating functional diversification among *Ep*RALPHs. *Ep*RALPH11 enhanced susceptibility to *E. pisi* in *Medicago truncatula*, localized to the nucleolus, and induced nucleolar fragmentation when heterologously expressed in *Nicotiana benthamiana* leaves. Its RNase domain exhibited T1 RNase activity in vitro, supporting the retention of a catalytically competent RNase domain and, together with its nucleolar localization, suggesting that *Ep*RALPH11 targets plant rRNA and disrupts nucleolar functions. The RNase domain induced cell death in *N. benthamiana*, whereas the full-length protein and catalytic mutants did not. Cell death induction required exclusive nucleolar localization of the RNase domain, and an extended N-terminal intrinsically disordered region suppressed this activity in the full-length protein. Together, our findings reveal a previously unrecognized mechanism regulating RNase activity in a dicot PM RALPH effector and provide new insights into the functional diversification of RALPHs and their adaptation to obligate biotrophy.

## Introduction

Powdery mildews (PM) are ascomycete fungi in the order Erysiphales, comprising ∼ 16 genera and 900 species and capable of infecting around 10,000 monocot and dicot plant species (1,2). The disease is easily recognized by the appearance of white, powdery symptoms on the aerial parts of plants, including leaves, stems, flowers, and pods. A distinguishing feature of PM fungi is their obligate biotrophy; they can colonize and complete their life cycle only on living plant tissues (3). For successful establishment on the host plant, PM fungi secrete an array of virulence proteins, termed effectors. The primary role of these effectors is to interfere with host metabolism and suppress immune signaling, creating a favorable environment for pathogen colonization (4). Pathogen effectors target diverse molecules within different compartments of plant cells and are classified as ‘virulent’ or ‘avirulent’ depending on whether they are perceived by the host immune system. ‘Virulent’ effectors evade host recognition and promote pathogen establishment, leading to effector-triggered susceptibility. ‘Avirulent’ effectors are (directly or indirectly) recognized by cognate intracellular plant resistance proteins, which leads to effector-triggered immunity (5). Investigations into PM effectors can, therefore, provide novel insights into fungal pathogenicity and host immunity.

PM effectors are primarily synthesized and secreted from haustoria, which are specialized fungal infection structures that develop within the apoplast of host epidermal cells (6,7). Early ‘omics’ studies defined PM effector candidates as proteins that harbor an N-terminal secretion signal, lack a transmembrane domain, and show limited sequence similarity outside the PMs (8,9). Using these sequence features, >490 candidate secreted effector proteins (CSEPs) were identified in the cereal PMs infecting barley (*B. graminis* f. sp. *hordei; Bgh*) and wheat (*B. graminis* f. sp*. tritici*; *Bgt*) (9,10). It is important to note that in subsequent studies on effector characterization, the same CSEPs were often assigned alternate and sometimes multiple names [e.g., Blumeria effector candidate (BEC), Bgt effector (BgtE), Bcg, AVR, etc.], which has made it challenging for researchers working in the field to follow the nomenclature. Nevertheless, structural analyses revealed that CSEPs harboring a ribonuclease domain and structurally resembling fungal T1 RNase are the most abundant, constituting ∼15% of the effector repertoire (9). These CSEPs were later named ‘RNase-like proteins associated with haustoria’ (RALPHs) as they were found to be predominantly expressed in haustoria (11). Despite harboring a ribonuclease domain, cereal PM RALPHs are pseudoenzymes, lacking the conserved catalytic residues required for ribonuclease activity (12,13). A comprehensive structural analysis of effector proteins from diverse fungal phytopathogens revealed that the RALPH effector family has undergone extensive expansion in the obligate biotrophic cereal PM *Bgh* (14). Subsequent structure-guided analysis further demonstrated that RALPH-related proteins constitute a major component of the secretomes of other *Blumeria* lineages and of several dicot-adapted PMs (15).

Functional characterization of cereal PM RALPHs has identified virulent (16) and avirulent members of this effector family (13,17,18). For example, host-induced gene silencing (HIGS) of two *Bgh* RALPHs, *BEC1011* and *BEC1054* (19), and three *Bgt* RALPHs, *SvrPm3^a1/f1^*, *Bcg6,* and *Bcg7* (20), was shown to reduce the ability of the fungi to penetrate and form haustoria in their respective host plants, indicating their role in virulence. A role in host immune modulation was ascribed to BEC1054, as it was found to interact with host proteins involved in defense and pathogen response, including glutathione S-transferase, malate dehydrogenase, and pathogenesis-related (PR) proteins PR5 and PR10 (16,21). In addition, transgenic expression of BEC1054 increased susceptibility to adapted pathogens in both monocot and dicot plants (16). The crystal structure of BEC1054 (PDB:6FMB) revealed that it is a close homolog of fungal T1 RNases and exhibits features suggestive of a role in nucleic acid binding. However, BEC1054 showed only weak binding specificity for plant ribosomal RNA (rRNA) in *in vitro* experiments. The study further showed that expression of BEC1054 partially inhibited the methyl jasmonate-induced cleavage of rRNA by plant ribosomal inhibiting proteins (RIPs), suggesting that this RALPH may bind to rRNA motifs recognized by plant RIPs. BEC1054-like RALPH effectors may, therefore, play an important role in promoting biotrophy by inhibiting the function of plant RIPs that would otherwise cause host cell death (16). A few cereal PM RALPHs are avirulent, as they are directly recognized by cognate host resistance proteins (17,18,22). For example, the *Bgh* RALPHs, AVR_A6_, AVR_A7_, and allelic AVR_A10_/AVR_A22_, are recognized by their cognate host resistance proteins MLA6, MLA7, MLA10, and MLA22 (12). However, these avirulent RALPHs lack RNase activity and exhibit only weak RNA binding. Overall, these studies suggest that RALPHs with RNA hydrolytic activity have not been retained during evolution in cereal PMs (13).

In contrast to cereal PMs, RALPH effectors from dicot-adapted PM fungi remain largely unexplored, with very limited functional characterization and experimental validation reported to date. *Erysiphe pisi (Ep)*, the primary causal agent of PM disease in pea, is an economically important pathogen that is responsible for severe yield losses in pea and other grain and forage legumes, including lentils, alfalfa, and *Medicago truncatula* (23,24). We previously identified 15 RALPH-like CSEPs from the *E. pisi* haustorial secretome through MCL, InterProScan, and/or BLAST analyses (25). HIGS of two *EpRALPHs*, *CSEP001*, and *CSEP009*, reduced fungal growth on pea leaves, indicating their role in pea PM virulence (25). Homology modeling revealed that CSEP001 and CSEP009 are structurally similar to fungal F1 ribonucleases, and, notably, the residues responsible for RNA hydrolysis are partially conserved in these two effectors. This led us to speculate that these *Ep*RALPHs may retain the ability to bind and degrade RNA, thereby ascribing distinct functions to this effector family in different PM lineages.

In the present study, we performed a comprehensive characterization of the RALPH effector repertoire of *E. pisi*. Structure-based comparative analysis of selected dicot-PM RALPHs identified multiple RNase-domain-containing RALPHs as a conserved feature of the *Erysiphe* lineage. Many *EpRALPH*s are preferentially expressed in haustoria and during the early stages of host colonization, suggesting roles in pathogen establishment. AlphaFold3-based structural analyses further showed that *Ep*RALPHs share a conserved RNase-like fold resembling fungal T1/F1 RNases but exhibit limited sequence and surface charge conservation, indicative of functional diversification. RALPH11/CSEP001 was identified as a putative RNA-binding protein that contains four of the six catalytic residues characteristic of T1/F1 RNases. Transient expression of RALPH11 in *Medicago truncatula* enhanced susceptibility to *E. pisi,* suggesting that this effector is required for pathogenesis on multiple legume hosts. RALPH11 localizes to the plant nucleolus and induces nucleolar fragmentation in *Nicotiana benthamiana,* suggesting that it may target rRNA and disrupt nucleolar functions. The RNase domain of *Ep*RALPH11 exhibits T1/F1 RNase activity *in vitro* and induces cell death upon transient expression in *N. benthamiana* leaves, whereas the full-length protein or catalytic mutants did not. Cell death also required exclusive nucleolar localization of the RNase domain, suggesting that its activity depends on rRNA binding and/or degradation. Our findings indicate that the RNase domain of RALPH11 is catalytically competent, whereas its activity is suppressed in the full-length protein by an N-terminal intrinsically disordered region. We propose that this regulatory mechanism enables RALPH11 to promote pathogenesis by interfering with host nucleolar functions while preventing host cell death, thereby supporting the pathogen’s obligate biotrophic lifestyle.

## Results

### Multiple RNase domain-containing RALPHs are restricted to the *Erysiphe* lineage

Studies on cereal PMs have shown that RALPH effectors are under extreme selection pressure to diversify their sequences to avoid host immune recognition (13,14). Therefore, in addition to sequence homology, we used 3D structural similarity-based approaches to identify RALPH effector candidates from the genomes of *Erysiphe pisi* (*Ep*; Palampur-1 isolate), and six additional dicot-adapted PM species, *E. necator* (*En)*, *E. pulchra (Epul)*, *Golovinomyces cichoracearum* (*Gc*), *G. orontii* (*Go*), *Oidium neolycopersici* (*On*), and *Parauncinula polyspora* (*Pp*), infectious on pea, grapevine, dogwood, Arabidopsis (*Gc* and *Go*), tomato, and Asian oak tree, respectively (**Fig. S1**; **Workbook S1**). We first predicted the secretomes of the respective PMs, and from these, candidate secreted effector proteins (CSEPs) were identified using EffectorP 3.0. We restricted our analysis to CSEPs because we were interested in identifying RALPHs that are also predicted to be effectors. We then used primary sequence analysis (InterProScan) and 3D structural similarity-based methods (ColabFold v1.5.5 and Rupee) to identify candidate RALPHs from the CSEPs (**Fig. S1; Workbook S1**).

In accordance with previous studies (15,26), we identified fewer RALPHs in *Ep* and other dicot-adapted PMs than reported for cereal PMs. *Go* has the largest secretome, with 483 members, followed by *Ep* with 345 (**Fig. 1A**). The secretomes of the remaining PMs are roughly similar, ranging from 223 to 252. A similar trend was observed for CSEPs, with *Go* (250) having the largest number among the dicot-adapted PMs, followed by *Ep* (189) and the others (70–115) (**Fig. 1A**). Notably, the *Erysiphe* species harbor a larger number of RALPHs than the other dicot lineages, with 22% of the *En*, 17% of the *Epul* and 18% of the *Ep* CSEPs classified as RALPHs (**Fig. 1A; Table S1**). In contrast, 1-12% of the CSEPs are RALPHs in the other dicot PM lineages, with *Go* containing the largest number. AlphaFold 3 (AF3) predicted structure analysis of RALPHs from these PMs revealed that most RALPHs contain a single ribonuclease domain, ranging in length from 80 to 130 amino acids (**Fig. 1B; Workbook S1**). However, a few RALPHs in the *Erysiphe* spp. contain 2, 3, 4, or 5 ribonuclease domains, with *Epul* harboring the maximum number of multi-RNase domain-containing RALPHs (**Fig. 1B**). Multi-RNase domain RALPHs are not present in the other dicot PMs, and as far as we know, have not been reported in cereal PMs.

**Figure 1.**
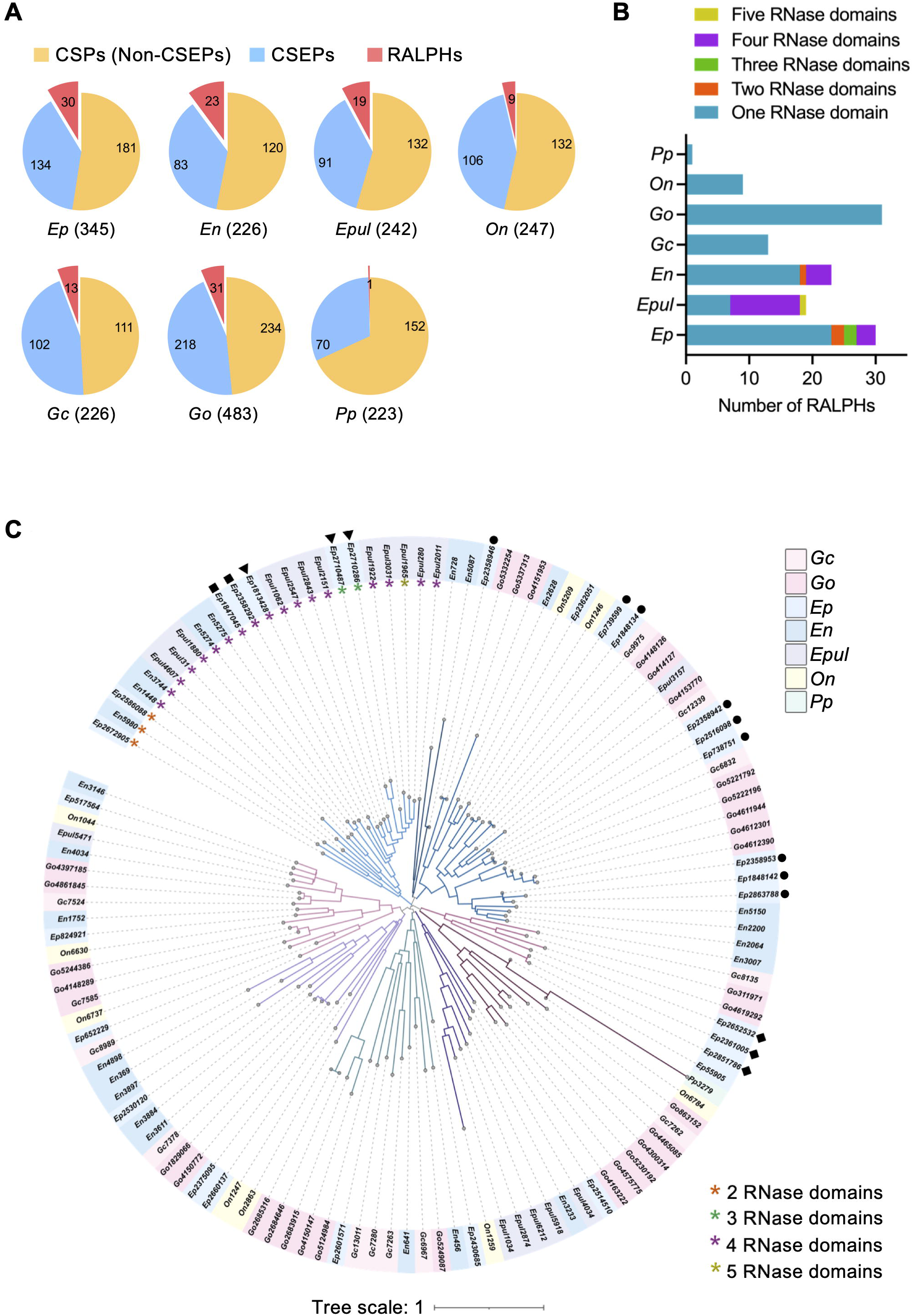
Genome-wide identification of PM RALPH effector candidates and their phylogenetic relationship. (A) Pie charts showing the distribution of non-CSEPs (CSPs that are not predicted to be CSEPs by EffectorP), CSEPs, and RALPH candidates in the secretomes of dicot-adapted PMs. The total number of CSPs in each secretome is shown in parentheses adjacent to the PM species name. (B) Graph showing the number of RALPHs with single and multiple RNase domains in each dicot-infecting PM based on AlphaFold3 models. (C) Structural relatedness of 126 dicot-PM RALPH effector candidates inferred using Foldtree. The scale bar indicates branch length. Branches are colored by group. PM RALPHs with more than one RNase domain analyzed through AF3 are indicated with asterisks. *Ep*RALPHs that form gene clusters on the genome are indicated with different shapes. Genes in the same cluster are marked with the same shape. *Ep, Erysiphe pisi; En, E. necator; Epul, E. pulchra; On, Oidium neolycopersici; Gc, Golovinomyces cichoracearum; Go, G. orontii; Pp, Parauncinula polyspora*

Given the low sequence identity and rapid diversification of PM effectors, traditional sequence-based phylogeny provided limited resolution (data not shown). Thus, a structural phylogeny using Foldtree (https://neurosnap.ai/service/Foldtree) was constructed to generate an alignment-free, distance-based tree and to compare the shared structural topologies and clade organization of the AF3-predicted structures of 126 dicot PM RALPH effector candidates. The resulting structure-based tree grouped the RALPHs into nine structural clusters (**Fig. 1C**), with members from different dicot PM lineages represented in all but two clusters. Notably, all multi-RNase-domain RALPHs were confined to these two clusters, suggesting a lineage-specific expansion of this subfamily in *Erysiphe*.

### *Ep* contains a higher number of RALPHs than previously estimated

For a more comprehensive analysis of the dicot PM RALPHs, we focused on the *Ep*RALPHs. In our previous paper describing the *Ep* haustorial transcriptome, we identified 15 RNase-like *Ep*CSEPs using InterPro, MCL family clustering, and BLAST analyses (25). To ascertain how many of these RNase-like CSEPs were also identified from the *Ep* genome, we performed a BLASTp analysis of the transcriptome-predicted CSEPs against the *Ep* Palampur-1 v2.0 protein database at MycoCosm. Of the 15 RNase-like CSEPs identified from the haustorial transcriptome, 5 showed >95% sequence identity at 100% query coverage with RALPHs identified from the *Ep* genome, 2 showed >98% sequence identity at partial (>60%) query coverage, whereas the remaining 8 showed moderate to low sequence similarity and low query coverage to RALPHs or other *Ep* proteins (**Table S2**). Therefore, genome mining identified 23 additional *Ep*RALPHs. Sequence-wise, RALPHs 3 and 17 are identical, and RALPHs 14 and 24 are identical except for an additional 8 amino acid residues near the C-terminal end of RALPH 24 (**Workbook S1**). We speculate that the incongruence between the genome and transcriptome data may, in part, be due to the presence of alternate splice forms or incorrectly assembled transcripts in the *Ep* haustorial transcriptome, which was largely *de novo* assembled using a fragmented and incomplete genome as a guide (25).

Phylogenetically related RALPHs in cereal PMs are often arranged tandemly in gene clusters (10). Therefore, we examined whether the phylogenetically related *Ep*RALPHs cluster in the genome. *RALPHs* located within 100 kb of each other on a scaffold were considered part of a cluster, as per (10). We found that 17 of the 30 *EpRALPHs* formed 4 clusters, with the smallest cluster containing 2 members and the largest cluster containing 9. Notably, the *EpRALPHs* that are physically clustered on the genome are also phylogenetically more closely related (**Fig. 1C**), indicating that local gene duplication may have contributed to the expansion of this family in the *Ep* genome, as observed for the cereal PMs (9).

### Many *EpRALPHs* display higher expression in haustoria and during early infection stages

In our previous study, eight of the nine RNase-like *CSEPs* tested by RT-qPCR showed preferential expression in haustoria compared with epiphytic tissues (25). For a more comprehensive analysis, we examined the spatial expression patterns of the *RALPHs* identified here, as well as those not assessed in our previous study. Transcripts were detected for all analyzed *EpRALPHs* except *RALPH08* and *12,* which are either expressed at very low levels or not under the conditions tested. Of the twenty-three *RALPHs* analyzed via RT-qPCR, fifteen showed significantly higher expression in haustoria compared to epiphytic mycelial tissues (**Fig. 2A**). Of these, except *RALPH28*, all showed ≥ 5-fold higher expression in haustoria compared to epiphytic mycelia. *RALPH11*, the highest expressed *CSEP (CSEP001)* in the *Ep* haustorial transcriptome (25), again showed the highest fold change in expression in haustoria versus epiphytic mycelia, confirming our previous results (**Fig. 2A**). *RALPH15* showed significantly lower expression in haustoria compared to epiphytic mycelia, whereas *RALPH01, 04, 16*, *19*, *20, 26 and 27* exhibited similar expression levels in haustoria and epiphytic mycelial tissues (**Fig. 2A**).

**Figure 2.**
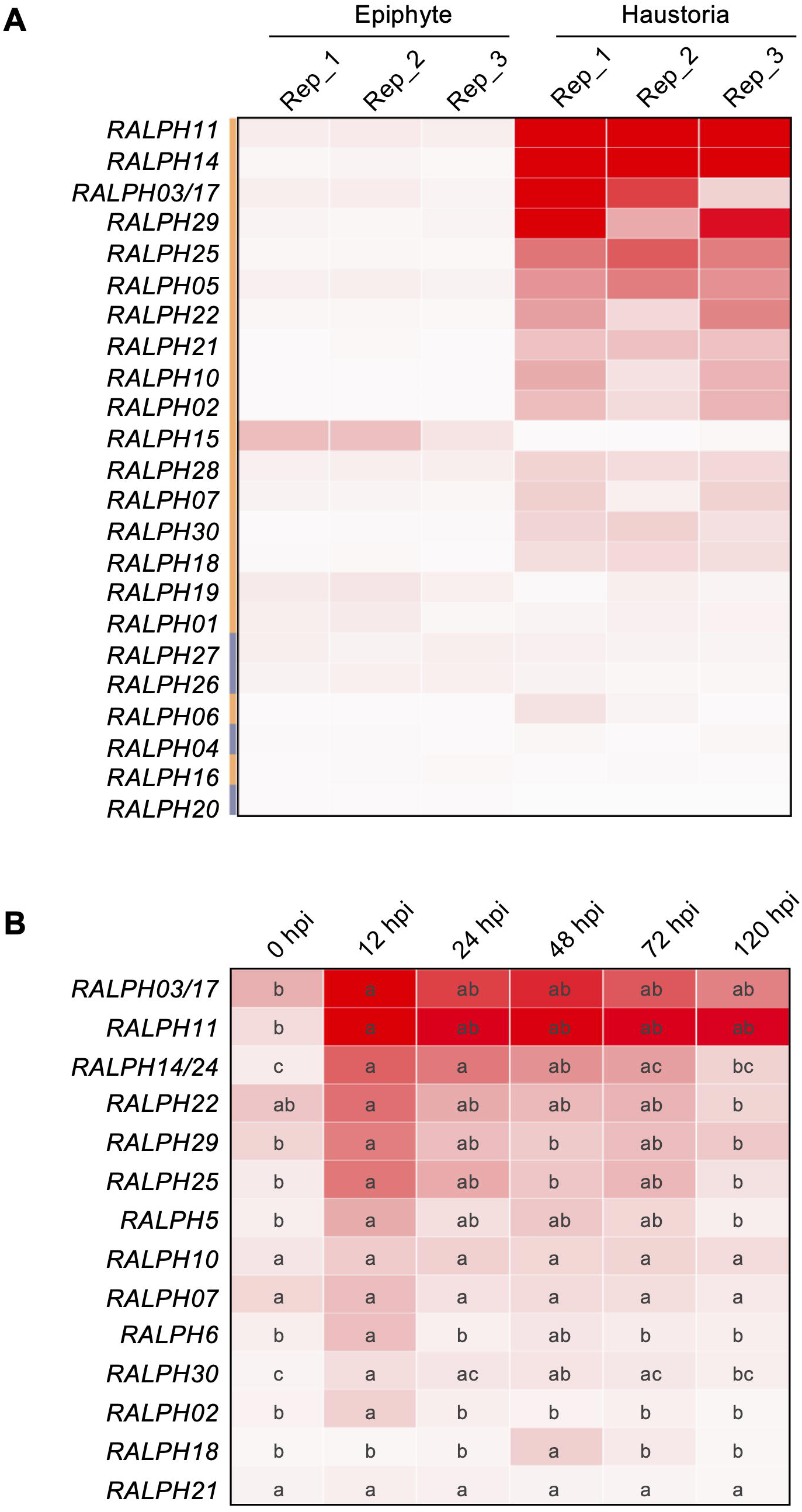
Spatial and temporal expression heatmaps of selected *EpRALPHs* analyzed via RT-qPCR. (A) Relative expression of *EpRALPHs* in epiphytic and haustorial tissues normalized to that of the reference genes *EpTUB2* and *Ep18S* rRNA from three independent biological replicate experiments. Statistical significance (p< 0.05) was computed via an unpaired t-test. The orange vertical bar indicates significant data, whereas the purple bar indicates non-significant data. (B) Relative expression (mean) of selected *EpRALPHs* at different time points after PM inoculation on pea leaves, normalized to that of the reference genes *EpTUB2* and *Ep18S* rRNA from three independent biological replicate experiments. hpi, hours post-inoculation. Statistical significance was computed via ordinary one-way ANOVA followed by Tukey’s multiple comparisons test. Statistically significant differences between time points are indicated via compact letter display. Heat maps were made using Clustergrammer (https://maayanlab.cloud/clustergrammer/) with rows ranked by sum.

The temporal expression patterns of the 15 *EpRALPHs* with ≥5-fold higher expression in haustoria than in epiphytic mycelia were analyzed at multiple time points after PM inoculation. Like *RALPH11*, the candidate effectors *RALPH02, 03/17, 05, 06, 14/24, 22, 25, 29* and *30* exhibited higher expression at the early infection stages (12-24 hpi) with peak expression at 12 hpi (**Fig. 2B**). These time points correspond to the fungal penetration and primary haustorium formation stages. Only *RALPH18* showed significantly higher expression at 48 hpi (**Fig. 2B**), corresponding to the colony expansion stage. The remaining three *RALPHs, 07, 10,* and *21,* showed similar expression levels across all infection time points tested (**Fig. 2B**). Overall, the temporal expression analysis suggests that many *RALPH*s are expressed early in infection and may facilitate host colonization.

### *Ep*RALPHs have a T1/F1 RNase-like structural scaffold

To gain structural insights into the RNase-like effector proteins, we analyzed the AlphaFold 3 (AF3) models of 29 unique *Ep*RALPHs (27). As RALPH14 and 17 are identical to RALPH24 and 3, respectively, only the latter were included in the structural analysis. The models of the full-length proteins reveal that the *Ep*RALPHs harbor a well-structured RNase-like scaffold with a high pLDDT (predicted Local Distance Difference Test) score (>50) along with a structurally disordered region with a very low pLDDT score (<50), either at the N-terminal (in most *Ep*RALPHs) or at the C-terminal (in some *Ep*RALPHs) (**Fig. 3; Fig. S2A**). Although most *Ep*RALPHs have a single compact RNase-like domain, RALPH18 and 21 contain two, RALPH22 and 23 contain three, and RALPH06, 07, and 10 have four RNase-like domains in tandem (**Fig. 3; Fig. S2B; Table S3)**. For identification, more than one domain within a RALPH is denoted by A, B, C, and D, each suffixed with its number. To confirm that the presence of multiple RNase domains in the seven *Ep*RALPHs is not an annotation error, we performed PCR amplification of their coding sequences using *Ep* cDNA as the template. In all cases, the CDS amplicon size obtained by PCR amplification of *Ep* cDNA closely matched the predicted CDS size reported in MycoCosm (**Fig. S3**). We then sequenced the amplicons and aligned them with the predicted gene sequences from MycoCosm using NCBI BLAST. The PCR amplicons of *EpRALPH06*, *10*, *18* and *21* showed 99-100% identity with their respective predicted CDS, matching the predicted exon-intron structure provided in MycoCosm (**Fig. S4**). By contrast, some or all predicted intron sequences were retained in the amplicons of *EpRALPH07*, *22*, and *23* (**Fig. S3B**; **S4**), indicating potential differences between predicted and actual transcripts and, consequently, proteins for these three *Ep*RALPHs.

**Figure 3.**
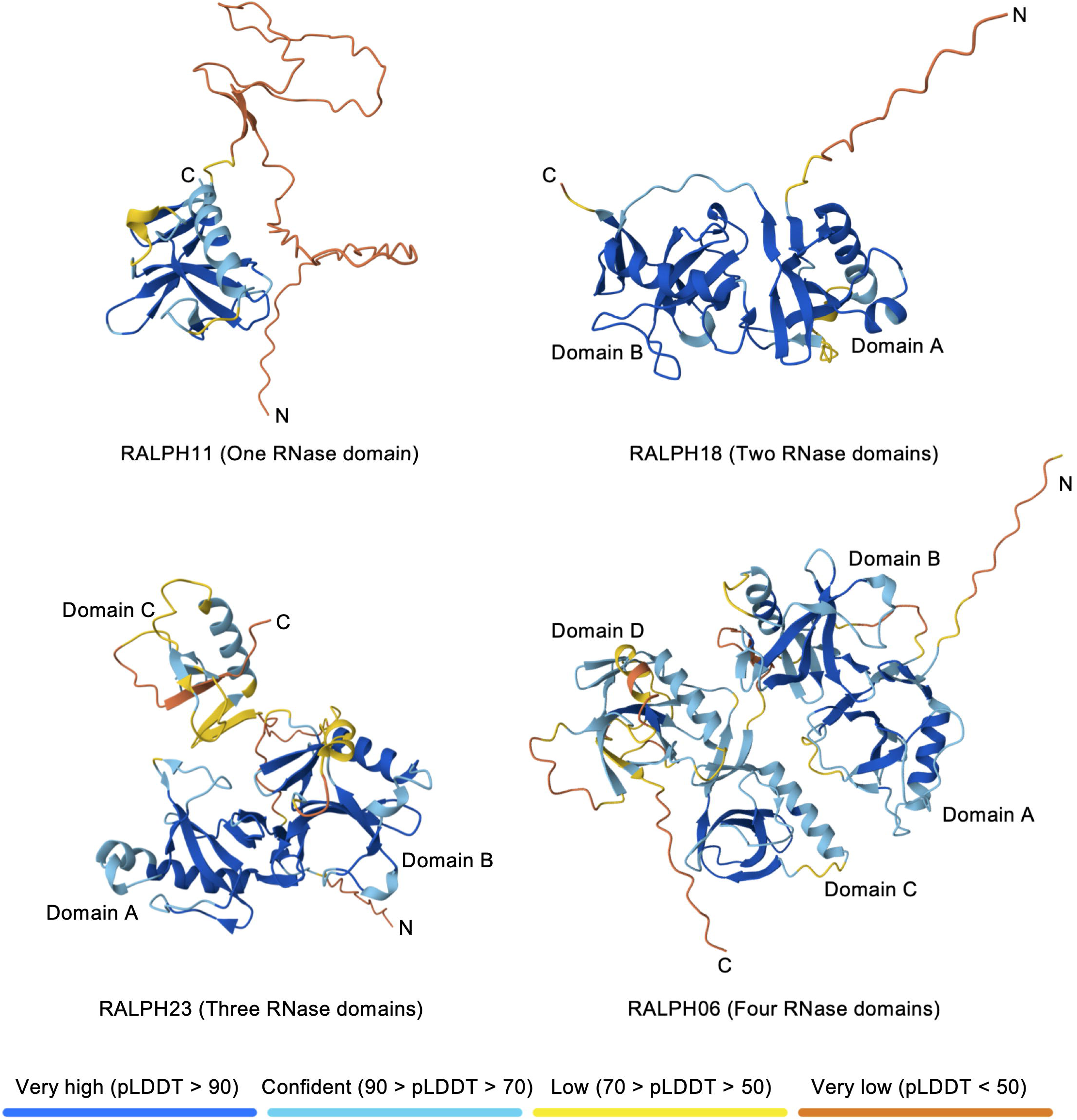
AlphaFold-3 predicted structures of representative single- and multi- RNase-domain full-length *Ep*RALPHs, colored based on the confidence score (pLDDT) from very low (orange), low (yellow), confident (light blue) to very high (dark blue), shown in the bar below. Lower pLDDT scores (orange) indicate the disordered stretch, and higher scores (blue) indicate greater confidence in the structure. The N- and C-termini of the RALPHs are labeled.

The AF3-predicted *Ep*RALPH ribonuclease domains were subjected to further analysis. The *Ep*RALPH RNase domains generally show a α/β-type structural fold typical of T1/F1 RNases, which generally consists of one peripheral β-sheet (consisting of 2 N-terminal β-strands) and one central β-sheet (consisting of 5 β-strands) packed on a long α-helix (28). The similarity of the *Ep*RALPH RNase domains with T1/F1 RNases was further assessed by structurally aligning each *Ep*RALPH RNase domain with the available structures of T1 (PDB:1RNT) and F1 (PDB:1FUT) RNases using FoldMason (**Fig. 4**). Each alignment was assigned an MSA LDDT (Multiple Sequence Alignment Local Distance Difference Test) score, which quantifies the degree of similarity (closer to 1 indicates greater similarity). The RNase domains of RALPH03, 11, 13, 18B, and 25 harbor a central β-sheet of 5 β-strands against a long N-terminal α-helix and are most similar to T1/F1 RNases (MSA LDDT score>0.8) (**Fig. 4; Table S4**). Remaining *Ep*RALPH RNase domains show a lower degree of structural similarity (MSA LDDT<0.8) to T1/F1 RNases due to the variations in the number of β-strands, observed in either the central or peripheral β-sheets, or in both, along with the presence of long loop-like structures and small helices between the β-strands (**Fig. 4; Table S4**). Overall, the structures of *Ep*RALPH RNase domains retain the core secondary structural elements of the canonical T1/F1 RNases; additional insertions contribute to structural differences, which may affect their function.

**Figure 4.**
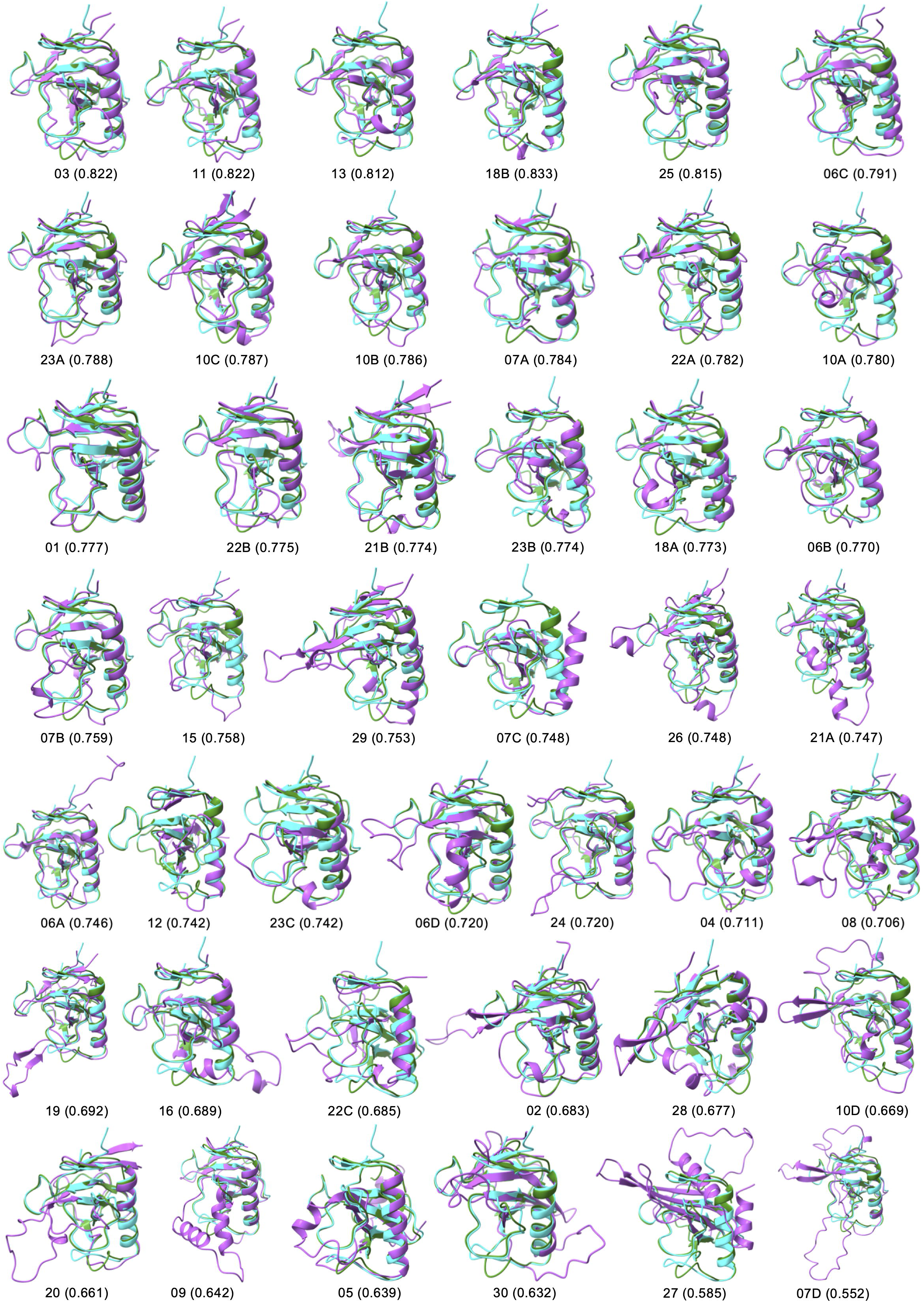
Structural alignment of ribonuclease domains of *Ep*RALPHs with the crystal structures of T1 (PDB:1RNT) and F1 (PDB:1FUT) RNases depicting structural similarity and conserved fold. The name of each *Ep*RALPH and MSA LDDT score (in parentheses) is depicted below the corresponding superimposition. The individual domains of the multidomain RALPHs are separately superimposed and denoted with suffixes A-D. T1 RNase, F1 RNase, and *Ep*RALPHs are colored in cyan, forest green and medium orchid, respectively.

Two characteristic disulfide bridges are hallmarks of T1 (Cys2-Cys10 and Cys6-Cys103) and F1 (Cys6-Cys103 and Cys24-Cys84) RNases. These disulfide bridges constrain the structure and provide stability to the protein (29). While one (Cys6-Cys103) of two disulfides is common to both RNases, the second is present within the peripheral β-sheet in T1 RNase (Cys2-Cys10) and within the central β-sheet in F1 RNase (Cys24-Cys84) (**Table S5**). The RNase domains of RALPHs 03, 04, 08, 09, 11, 13, 15, 16, 20, 24, and 25 each contain two disulfides at the canonical positions as in F1 RNase; others show pronounced variation, indicating differences in the stability of these proteins. For example, in addition to the two canonical disulfide bonds, the RNase domains of RALPHs 02, 19, 26, 28, and 30 have an extra disulfide bond at positions other than in T1/F1 RNases. The RNase domains of RALPH05 and 12 do not harbor any disulfide bond, possibly due to the lack of the peripheral β-sheet and a complete central β-sheet (2 β-strands instead of 5 β-strands), respectively (**Tables S4 & S5)**. Multidomain RALPHs have only one domain that possesses both the disulfides conserved in F1 RNase, while the other domains have the common T1/F1 RNase disulfide or no disulfides at all (**Table S5**). Thus, structural investigation revealed that most *Ep*RALPHs have a core resembling that of T1/F1 RNases, with 21 RALPHs having at least one domain with two pairs of Cys residues, as found in F1 RNase, indicating that *Ep*RALPHs are more related to F1 than to T1 RNase.

### *Ep*RALPHs share limited sequence similarity

The multiple sequence alignment shows that the *Ep*RALPHs exhibit limited sequence conservation among themselves, with conservation restricted to only a few hydrophobic residues in the core of each molecule, despite a common structural fold (**Fig. S5**). Despite the limited sequence similarity in the primary protein sequence, the identification of a single intron at the same relative position in several *Bgh* and a few *Bgt* RALPHs had previously led to the conclusion that RALPHs have evolved from a single RNase or RNase-like ancestor (9,11). To determine whether this feature is conserved in the single-intron *EpRALPHs*, we analyzed intron positions in the multiple-sequence alignment. We found that in thirteen out of sixteen single intron-containing *EpRALPH*s (**Fig. S6A**), the intron is in the same relative position as found previously in the *Bgh* RALPHs (**Fig. S6B**), indicating a common phylogenetic origin of cereal- and dicot-adapted PM RALPHs.

### *Ep*RALPH11 contains four putative T1/F1 RNase catalytic residues

The canonical fungal T1/F1 RNases cleave RNA at the 3’ end of the guanosyl residue and contain 6 catalytic residues, including Tyr38 (Y38), His40 (H40), Glu58 (E58), Arg77/76 (R77/76), His92/91, (H92/91) and Phe100/99 (F100/99) (30–32). Sequence alignment of the RNase domains shows that no *Ep*RALPH has all six catalytic residues present (**Fig. S5**). The Tyr residue corresponding to Y38 of T1/F1 enzyme is present in at least one RNase domain of 12 RALPHs. The first and second His residues corresponding to T1/F1 enzyme H40 and H92/H91, respectively, are not present in any of the RALPHs; however, RALPHs 03, 11, 12, and 22C contain a His at position 39, corresponding to position 42 in T1/F1 RNases, and RALPHs 3 and 11 contain a His corresponding to position 93/92 in the T1/F1 RNases. The Glu corresponding to E58 of T1/F1 enzyme is present in 6 RALPHs, including RALPHs 03 and 11. The Arg corresponding to R77/76 in T1/F1 enzyme is present in at least one RNase domain of 18 RALPHs. The Phe corresponding to F100/99 of T1/F1 enzyme is present in RALPH01, 6B, 16, and 29. Hence, *Ep*RALPH3 and 11, which show high sequence identity (∼86%) between them, contain three putative catalytic residues, E52, R71, and H87, within their RNase domains (**Fig. S5**) and are likely to be active proteins. This is consistent with our previous analysis, where the same three catalytic residues were predicted in CSEP001 (RALPH11) based on sequence alignment with T1/F1 RNases (25). However, the other RALPHs that lack one or more of these residues are likely to be deficient in RNase activity.

To determine whether *Ep*RALPHs bind RNA, their RNase domain sequences were submitted to I-TASSER, and the predicted ligand, binding-site residues, and associated confidence scores (C-scores) were analyzed. As a test case, the T1 RNase structure produced a C-score of 0.99 with the ligand 2’-GMP (Guanosine-2’-monophosphate), with all 6 catalytic residues identified as ligand-binding site residues, which is consistent with the crystal structure (PDB:1RNT) (**Fig. 5A; Table S6**). Although 2’-GMP was predicted as the top ligand for most *Ep*RALPHs, only three were high-confidence predictions (C-score >0.75), with RALPH11 having the highest C-score (0.95), followed by RALPH13 (0.87) and RALPH21B (0.85) (**Table S6**). Two (E52, R71) catalytic residues identified via sequence alignment (**Fig. S5**) were predicted as 2’-GMP-binding residues in RALPH11, two (E50 and R69) in *Ep*RALPH13, and none in RALPH21B (**Table S6**). I-TASSER also predicted R86, corresponding to H92/91 in T1/F1 RNases, as a ligand-binding residue in RALPH11 and in several other *Ep*RALPHs (**Table S6**; **Fig. 5B**). In addition, H39 corresponding to position 42 in T1/F1 RNases was predicted as a ligand-binding residue in RALPH11 (**Table S6; Fig. 5B**). Structure-based sequence alignment of T1 RNase (PDB: 1RNT) and the I-TASSER-predicted structure of 2’-GMP-bound *Ep*RALPH11 (**Fig. 5C**) verified the sequence-based alignment. Hence, based on sequence alignment and I-TASSER prediction, we identified H39, E52, R71 and R86 as putative catalytic residues in *Ep*RALPH11.

**Figure 5.**
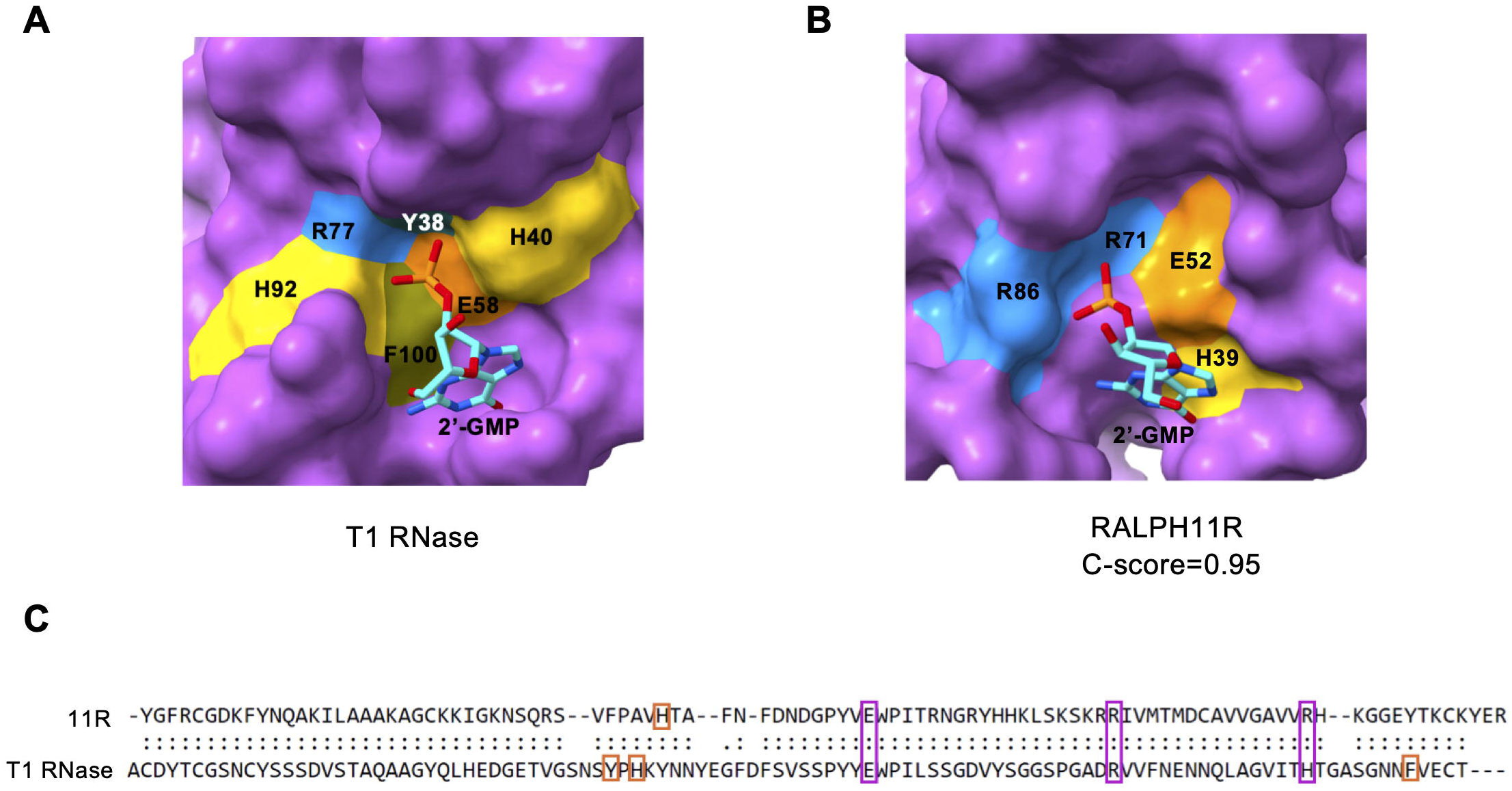
(A) Structures of 2’-GMP-bound T1 RNase (PDB: 1RNT) and (B) I-TASSER predicted *Ep*RALPH11R with 2’-GMP indicating active site residues. The protein is shown in surface representation with Tyrosine, Arginine, Glutamate, Histidine, Phenylalanine, and 2’-GMP in teal, dodger blue, dark orange, gold, olive and cyan, respectively. (C) TM-Align result showing the positions of active-site residues in RALPH11R relative to T1 RNase.

Next, we examined the electrostatic surface potential of *Ep*RALPHs. *Ep*RALPHs analyzed in this study differ significantly from classical T1/F1 RNases in their surface charge distributions. While the classical RNases have fewer basic residues, most *Ep*RALPHs, including RALPH11, which harbors four catalytic residues, show a highly positively charged surface (**Fig. S7**). The presence of positively charged residues indicates a strong binding affinity for negatively charged RNA, suggesting that *Ep*RALPHs that lack the conserved catalytic residues may perform RNA-related functions within the host beyond RNA degradation.

Overall, a comparison of the structure and charge distribution reveals that *Ep*RALPHs might have originated from a common ancestor and, during evolution, have accumulated sequence variations while retaining the common structural fold, thereby facilitating the diversification of RALPH function. Since the structural analysis indicated that *Ep*RALPH11 may have a high propensity to bind and degrade RNA, we selected this RALPH for further functional characterization.

### *Ep*RALPH11 transient overexpression enhances PM susceptibility of *Medicago truncatula*

We previously showed that host-induced gene silencing (HIGS) of *EpRALPH11* (*CSEP001*) reduces PM virulence on a highly susceptible pea genotype (25). To explore whether RALPH11 overexpression enhances host susceptibility to *E. pisi*, we transiently expressed a codon-optimized version of RALPH11 (lacking the N-terminal signal peptide) in the moderately susceptible *Medicago truncatula* R108 genotype. Medicago R108 leaves were infiltrated with GFP-RALPH11 or empty vector (negative control), and 2 days post-infiltration, leaves were inoculated with *E. pisi,* followed by fungal growth stage quantification at 48 hpi (hours post-inoculation). At 48 hpi, 38% of the conidia were in the secondary hyphal stage in GFP-RALPH11-infiltrated leaves, compared to 12% in empty vector controls, where appressorial stage conidia were predominantly higher in number (**Fig. 6A-B**), indicating that *Ep*RALPH11 overexpression enhances PM susceptibility of Medicago. The presence of GFP-RALPH11 and GFP proteins in the infiltrated leaves was confirmed via Western blotting at 3 days post-infiltration (**Fig. 6C**). These findings, combined with our previous HIGS results (25), suggest that RALPH11 is critical for *E. pisi* virulence on multiple legume hosts (pea and Medicago).

**Figure 6.**
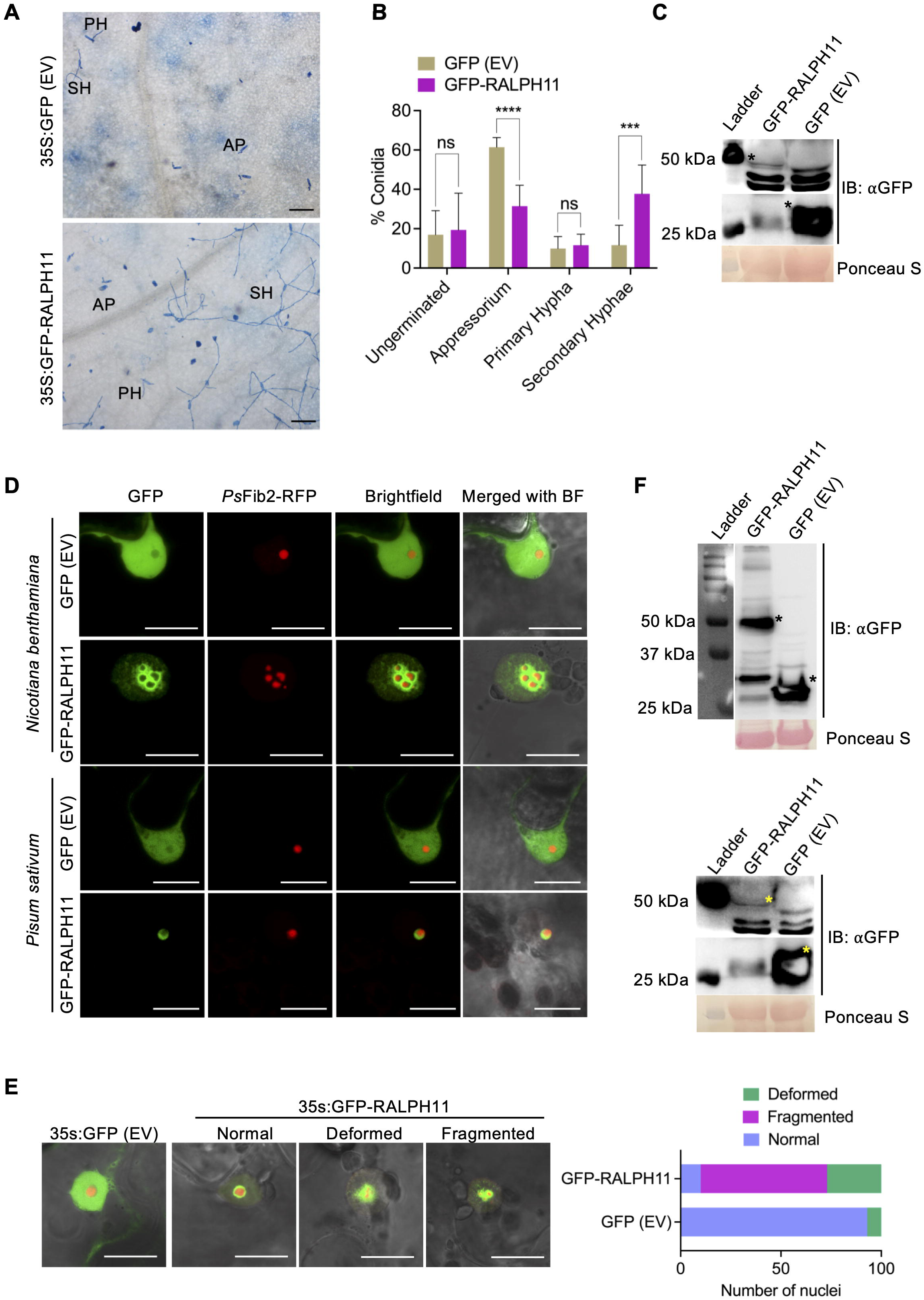
Transient expression of *Ep*RALPH11 enhances PM susceptibility of *Medicago truncatula* and triggers nucleolar fragmentation in *N. benthamiana*. (A) Representative images showing *E. pisi* growth on GFP (Empty vector, EV)- and GFP-RALPH11-infiltrated *M. truncatula* R108 leaves at 48 hpi. Scale bar, 200 µm. AP, appressorium; PH, primary hypha; SH, secondary hyphae (B) The mean percentage (± SD) of *E. pisi* conidia that reached different infection stages on GFP and GFP-RALPH11-infiltrated Medicago leaves at 48 hpi assessed from 2000 conidia on a total of 8 leaflets from two independent experiments. Significant differences were computed using an unpaired t-test (***p ≤ 0.001; ****p ≤ 0.0001; ns, not-significant). (C) Anti-GFP immunoblot displaying GFP-RALPH11 and GFP proteins (indicated by asterisks) in extracts of Medicago leaves at 3 days post-infiltration of the respective constructs. Ponceau S staining of the PVDF membrane shows protein loading. (D) Representative confocal images showing localization of GFP-*Ep*RALPH11 or GFP (EV) upon transient expression in *N. benthamiana* (above) and *P. sativum* (below). *Ps*Fib2-RFP was used as a nucleolus marker. (E) Representative confocal images showing nucleolar morphology phenotypes in GFP-*Ep*RALPH11- or GFP (EV)-infiltrated *N. benthamiana* leaves at 2 days-post infiltration; Scale bar, 15 µm (F) Anti-GFP immunoblot displaying GFP-RALPH11 and GFP proteins (indicated by asterisks) in extracts of *N. benthamiana* (above) and *P. sativum* (below) leaves at 3 days post-infiltration of the respective constructs. RuBisCO stained by Ponceau S was used as a total protein loading control.

### *Ep*RALPH11 localizes to the plant nucleolus and induces nucleolar fragmentation in *Nicotiana benthamiana*

We previously showed that *Ep*RALPH11 (CSEP001) localizes to the nucleus when transiently expressed in *N. benthamiana* (25). Using the Nucleolar Localization Sequence Detector tool (https://www.compbio.dundee.ac.uk/nod/index.jsp), we identified a nucleolar localization signal (NoLS) in RALPH11 between amino acid positions 62-110. To establish whether RALPH11 specifically localizes to the nucleolar sub-compartment, the codon-modified version of its coding sequence (minus SP) was tagged with GFP at its N-terminus to generate the 35S promoter-driven GFP-RALPH11 construct. The GFP-RALPH11 construct was co-infiltrated with the nucleolus marker *Ps*Fib2-RFP (*Pisum sativum* homolog of *At*Fibrillarin2, tagged with RFP) into the abaxial side of *N. benthamiana* (non-host) and *Pisum sativum* (host) leaves and imaged via confocal microscopy at 2- and 3-days post-infiltration, respectively. In contrast to GFP’s nucleocytoplasmic localization in empty vector-infiltrated leaves, GFP signals were restricted to the nucleus and mainly concentrated at the nucleolar rim in GFP-RALPH11-infiltrated *N. benthamiana* and *P. sativum* leaves (**Fig. 6D**). Faint GFP signals were also visible throughout the nucleus in *N. benthamiana,* but not in *P. sativum* leaves (**Fig. 6D**). Intriguingly, unlike in *P. sativum*, GFP-RALPH11 overexpression affected nucleolar morphology in *N. benthamiana* leaves. Nearly 90% of the nucleoli in GFP-RALPH11-infiltrated leaves appeared deformed or fragmented, with the nucleolar protein Fibrillarin2 localized to multiple discrete foci rather than a single nucleolar focus. In contrast, only 7% of nucleoli in empty vector-infiltrated leaves displayed this phenotype, with *Ps*Fib2 predominantly localized to a single nucleolar site (**Fig. 6E)**. We speculate that differences in RALPH expression levels may partially account for the distinct nucleolar phenotypes in the two plant systems, as higher GFP-RALPH11 protein accumulation was detected in *N. benthamiana* than in *P. sativum* leaves (**Fig. 6F**). The nucleolar localization of RALPH11 suggests that it may interact with rRNA and interfere with the normal functioning of this sub-nuclear compartment.

### Recombinant *Ep*RALPH11 protein exhibits T1 RNase activity

Structural analysis revealed that *Ep*RALPH11 may be catalytically active, as a few residues responsible for T1/F1 RNase activity are present in this effector (**Fig. 5**). To experimentally validate this prediction, we expressed and purified *Ep*RALPH11’s RNase domain (RALPH11R) in *E. coli* (**Fig. S8A**) and performed an *in vitro* RNase activity assay on total RNA isolated from pea leaves. Attempts to express and purify the full-length protein (RALPH11F) were unsuccessful, likely due to the structurally disordered N-terminal region. Incubation with T1 RNase (positive control) for 60 min resulted in complete degradation of pea RNA with no RNA Integrity Number (RIN) assigned to the sample (**Fig. 7A**). Treatment with GST-RALPH11R resulted in near complete degradation of pea RNA with a RIN of 2.9 (**Fig. 7A)**. In both cases, the 25S and 18S rRNA peaks were not visible or were diminished in the electropherograms. In comparison, the RIN value of pea RNA incubated with GST-RALPH02R (an *Ep*RALPH lacking all T1 RNase catalytic residues, thereby serving as a negative control) was 6.2, with the 25S and 18S rRNA peaks visible in the electropherogram. Similarly, the RIN value of the RNA samples treated with GST alone or nuclease-free water (no protein) was 8.8, with intact 25S and 18S rRNA peaks, indicating that the integrity of the RNA was largely unaffected in these samples (**Fig. 7A**).

**Figure 7.**
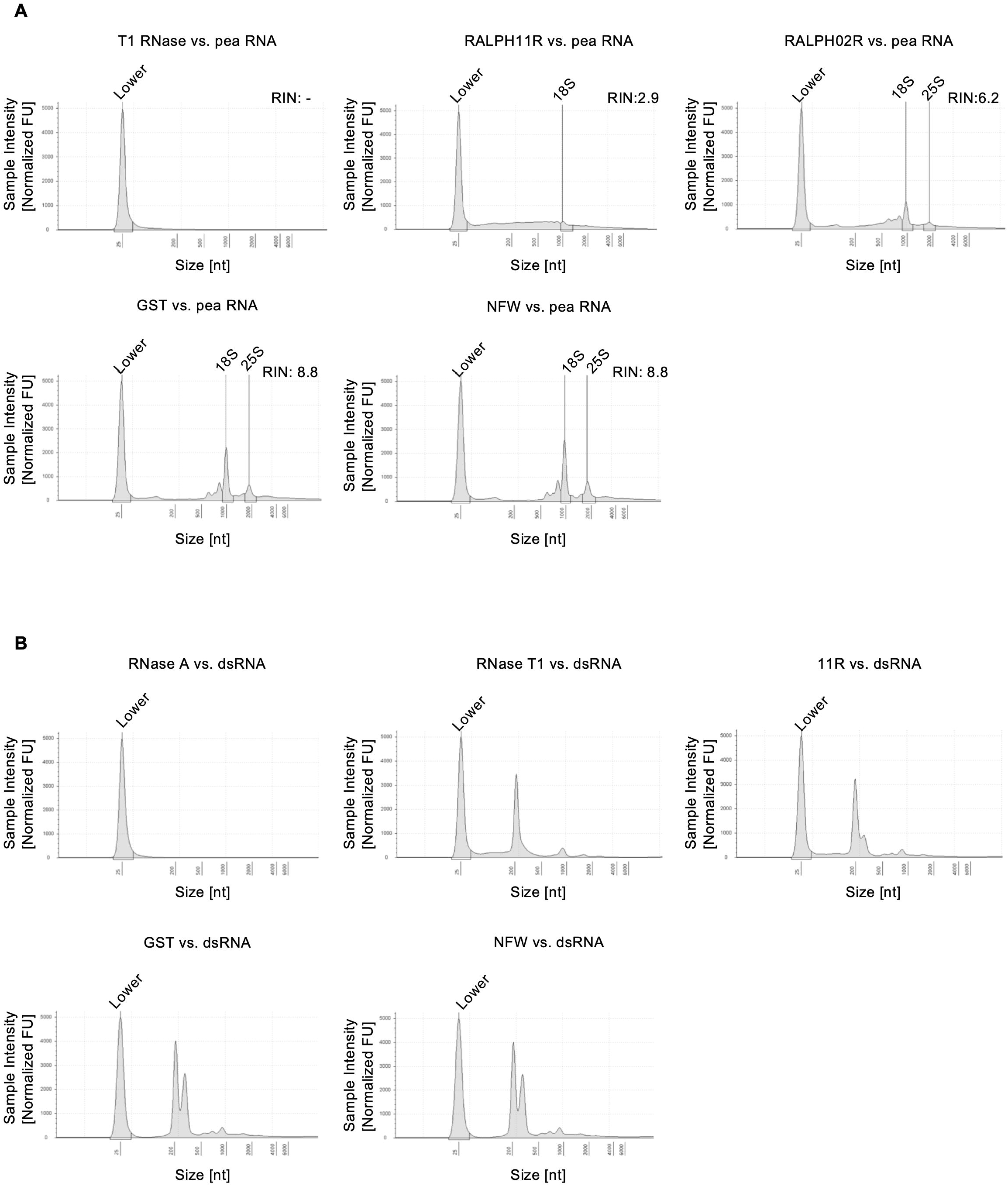
Recombinant *Ep*RALPH11R protein exhibits T1/F1 RNase activity. (A) Electropherograms showing the integrity of pea total RNA incubated with RNase T1, GST-*Ep*RALPH11R protein, GST-*Ep*RALPH02R protein, GST protein, or nuclease-free water (NFW), analyzed on an Agilent TapeStation. RIN, RNA Integrity Number. A higher RIN value indicates better RNA quality and integrity. (B) Electropherograms showing the integrity of GFP dsRNA incubated with RNase A, RNase T1, GST-*Ep*RALPH11R protein, GST, or NFW, analyzed on an Agilent TapeStation.

The T1/F1 family of RNases is known to cleave single-stranded RNA (ssRNA) but not double-stranded RNA (dsRNA) (33,34). To demonstrate that RALPH11R harbors a T1/F1 RNase-like activity, we performed the RNase activity assay using *in vitro-*transcribed GFP dsRNA as the RNA substrate. Treatment with RNase A, which can cleave both ssRNA and dsRNA (33), resulted in complete degradation of the dsRNA (**Fig. 7B**). In contrast, the GFP dsRNA remained intact when treated with GST-RALPH11R, T1 RNase, GST and nuclease-free water (no protein) under the same reaction conditions (**Fig. 7B**). The *in vitro* RNase activity against ssRNA (pea RNA) included in this experiment, showed similar results like the previous experiment, with complete RNA degradation by RNase A and RNase T1 (No RIN), RIN 1.8 with GST-RALPH11R, and no degradation by GST and nuclease-free water (**Fig. S8B).** The specificity for ssRNA indicates that RALPH11R possesses T1/F1-like RNase activity.

### *Ep*RALPH11R overexpression induces cell death in *N. benthamiana* in a nucleolar localization-dependent manner

Ribonuclease effectors secreted by many hemibiotrophic fungal pathogens induce plant cell death when transiently overexpressed in *N. benthamiana*, a feature that is dependent on their RNase activity [e.g., (35–37)]. To test whether the catalytically active *Ep*RALPH11R induces cell death, RALPH11R and RALPH11F were transiently overexpressed in *N. benthamiana*. The RNase domain of a previously characterized *Fusarium oxysporum* RNase effector [*Fo*SRE1, homologous to *St*SRE1; (36)] was used as a positive control. RALPH02 RNase domain (RALPH02R) and full-length (RALPH02F) proteins were used as negative controls. All proteins used in this study were tagged with GFP at the N-terminus and agroinfiltrated into *N. benthamiana* leaves, and their expression was confirmed by western blotting (**Fig. 8A**). At 5 days post-infiltration, cell death was observed in *Ep*RALPH11R and *Fo*SRE1R-infiltrated leaves, but not in RALPH11F, RALPH02F, 02R, or empty vector-infiltrated leaves (**Fig. 8A; Fig. S9A**). This indicates that the RALPH11F RNase domain, rather than the full-length protein, induces cell death in *N. benthamiana* leaves. Overexpression of RALPH11F did not induce cell death in *P. sativum* leaves either (**Fig. S9E**).

**Figure 8.**
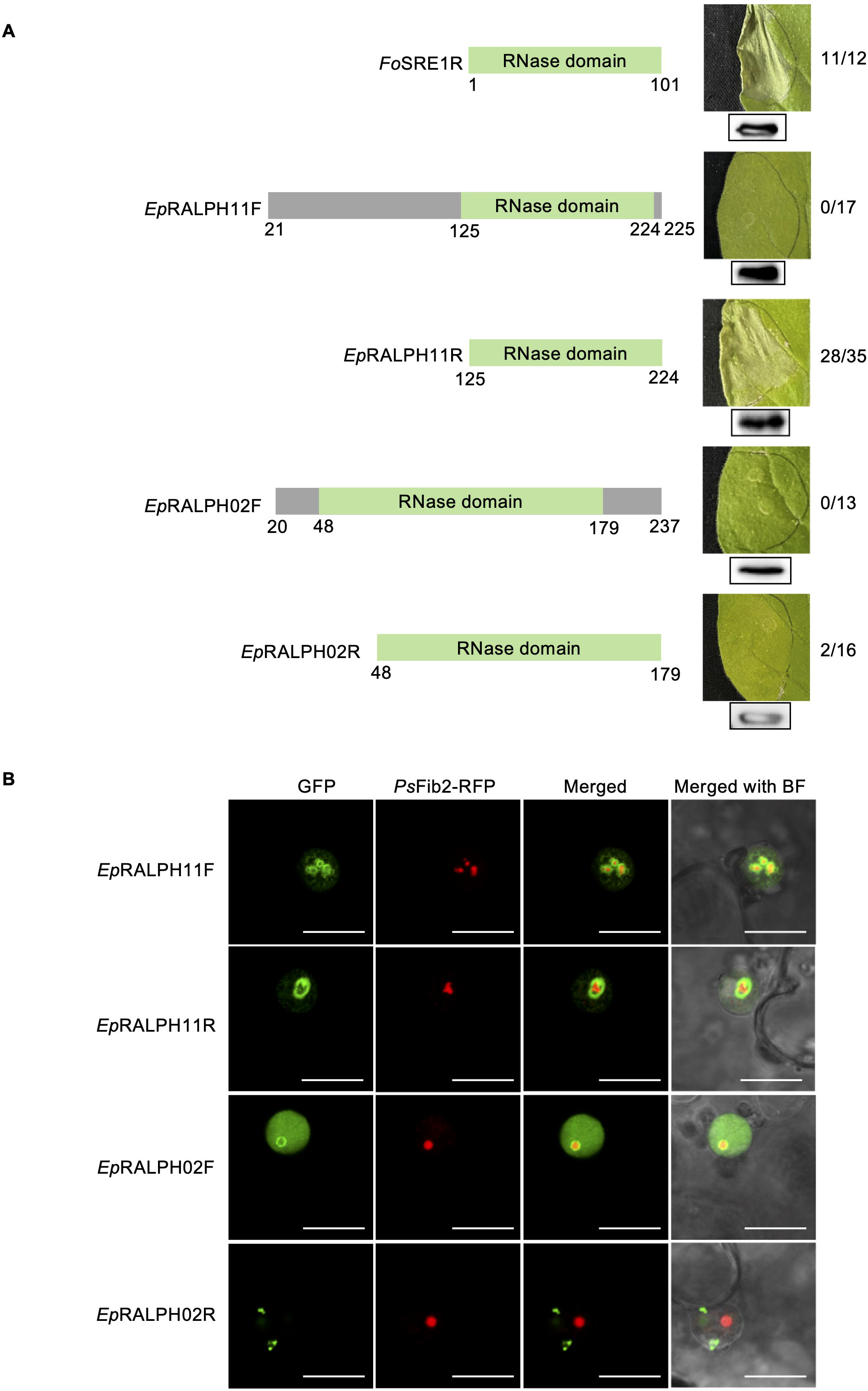
*Ep*RALPH11R induces cell death in *N. benthamiana*. (A) Left panel: Domain architecture of *Fo*SRE1 (positive control) and the *Ep*RALPHs transiently expressed in *N. benthamiana* leaves; Right panel: Corresponding representative leaf images showing presence or absence of cell death symptoms at 5 days post-infiltration of the respective constructs; F, full-length; R, RNase domain. Shown next to each leaf is the number of infiltrated leaves showing cell death symptoms/total number of infiltrated leaves. Displayed below each leaf image is the immunoblot confirming protein expression in *N. benthamiana* leaves transiently expressing the indicated proteins, using anti-GFP antibodies. RuBisCO stained by Ponceau S was used as a total protein loading control. (B) Representative confocal images showing the nuclear/nucleolar localization pattern of the indicated proteins. *Ps*Fib2-RFP was used as the nucleolus marker; Scale bar, 15 µm.

Nuclear localization of RNase effectors secreted by hemibiotrophic fungal pathogens is essential to induce plant cell death (35,36). To determine whether the nuclear localization pattern of the RALPH proteins affects their cell death-inducing ability, the *in-planta* localization of each protein was assessed at 2 days post-infiltration using confocal microscopy. GFP-RALPH11F signals were dispersed in the nucleus as well as aggregated around fragmented nucleoli, as observed previously; however, GFP*-*RALPH11R signals were visible only at the nucleolus (**Fig. 8B)**. In contrast, GFP-RALPH02F signals were mainly dispersed throughout the nucleus, and RALPH02R showed punctate signals within the nucleus but not at the nucleolus (**Fig. 8B)**. Notably, except GFP-RALPH11F, none of the proteins induced nucleolar fragmentation. Taken together, our findings suggest that nucleolar localization may be required for RALPH11R’s ability to induce cell death in *N. benthamiana*.

### *Ep*RALPH11R-induced cell death depends on the presence of catalytic residues

To determine whether cell death induced by RALPH11R overexpression is dependent on RNA degradation, four putative catalytic residues - H39, E52, R71 and R86 - were mutated to Alanine, individually and in combination, by site-directed mutagenesis (SDM), and the mutated GFP-RALPH11R constructs were agroinfiltrated into *N. benthamiana* leaves. Protein expression was confirmed by western blotting at 2 days post-infiltration (**Fig. 9A**). Like wildtype RALPH11R (RALPH11R^WT^), RALPH11R^E52A^ and RALPH11R^R71A^ mutants induced cell death at 5 days post-infiltration (**Fig. 9A**; **Fig. S9B-C**). In contrast, mutation at H39 or R86 (RALPH11R^H39A^ and RALPH11R^R86A^) abrogated RALPH11R’s cell death-inducing activity. The quadruple mutant (RALPH11R^HERR-A^), in which all 4 catalytic residues were mutated, also did not trigger cell death (**Fig. 9A**; **Fig. S9B-C)**. Taken together, these findings suggest that two of the four putative catalytic residues (H39 and R86) are critical for the cell death-inducing activity of RALPH11R in *N. benthamiana.* Notably, R86, which occupies the position corresponding to the canonical T1/F1 RNase H92, may participate in RNA binding and/or catalysis in RALPH11R.

**Figure 9.**
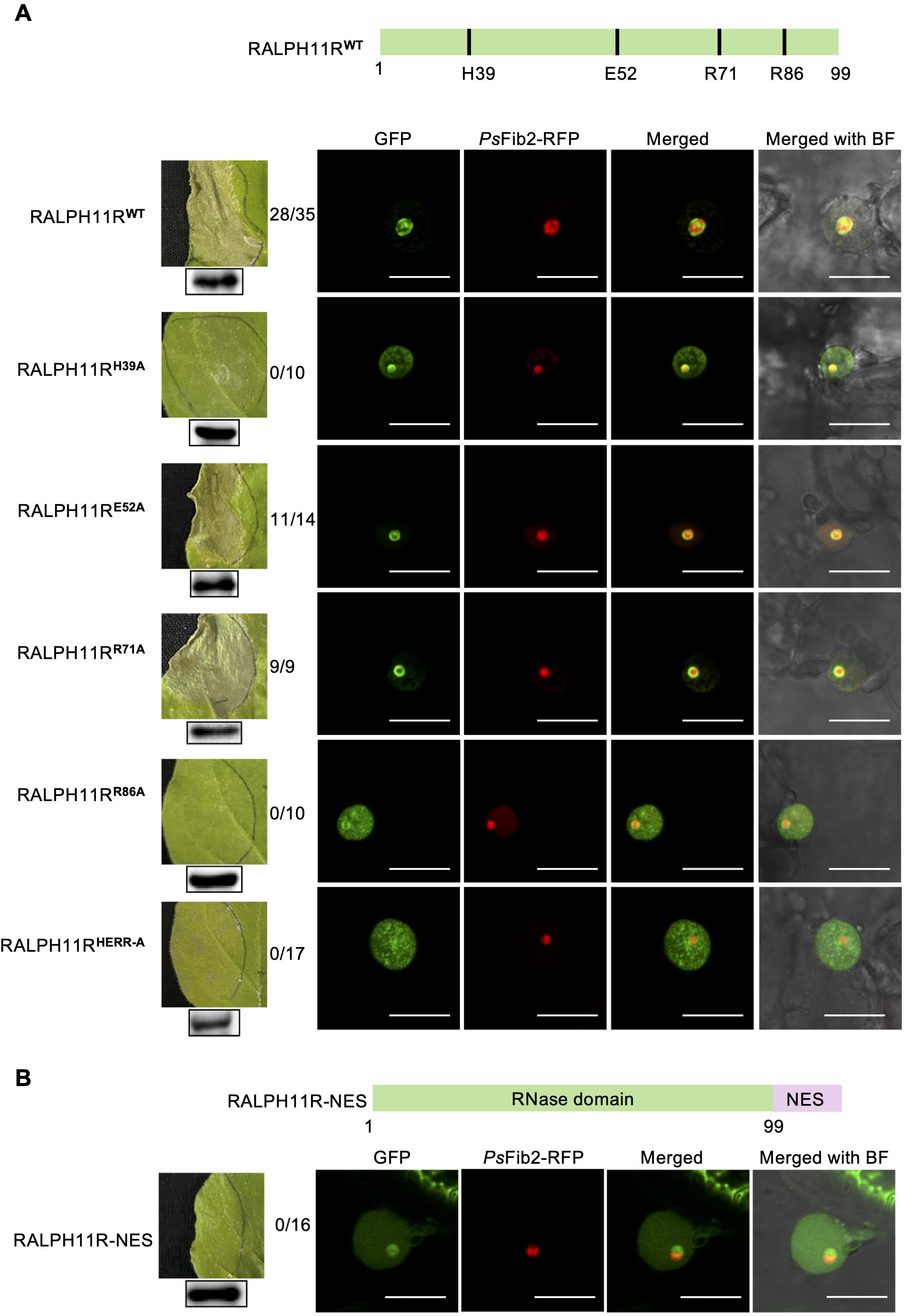
Cell-death-inducing activity and nucleolar localization analyses of the putative catalytic residue mutations in *Ep*RALPH11R. Left: *N. benthamiana* leaves were examined 5 days after transient expression of (A) *Ep*RALPH11R mutants and (B) *Ep*RALPH11R tagged with an NES (Nuclear Export Signal). The construct expressing the wildtype *Ep*RALPH11R was used as the positive control. Shown next to each leaf is the number of infiltrated leaves showing cell death symptoms/total number of infiltrated leaves. Immunoblotting results of expressed proteins at 2 days post-infiltration are shown below the leaves. Right: Representative confocal images showing the nuclear/nucleolar localization pattern of the indicated proteins. *Ps*Fib2-RFP was used as the nucleolus marker; Scale bar, 15 µm.

To assess whether nucleolar localization is affected in the different mutants, confocal microscopy was performed on *N. benthamiana* leaves at 2 days post-infiltration. Of the 4 mutants, RALPH11R^E52A^ and RALPH11R^R71A^, which caused cell death, showed nucleolar localization like RALPH11R^WT^ (**Fig. 9A**). In contrast, GFP signals of RALPH11R^H39A^, RALPH11R^R86A^, and the quadruple mutant RALPH11R^HERR-A^, which did not cause cell death upon expression, were more dispersed throughout the nucleus and less aggregated around the nucleolus (**Fig. 9A**). From these observations, we speculate that specific localization of RALPH11R at or around the plant nucleolus plays an important role in the induction of cell death. To further support this conclusion, GFP-RALPH11R was tagged with a Nuclear Export Signal [NES; (35)] and agroinfiltrated into *N. benthamiana* leaves. Confocal microscopy analysis revealed a dispersed nuclear localization pattern for GFP-RALPH11R-NES, as in RALPH11R^H39A^, RALPH11R^R86A^ and RALPH11R^HERR-A^, along with cytoplasmic localization (not observed in the above mutants) at 2 days post-infiltration (**Fig. 9B**). Accordingly, cell death was not induced in GFP-RALPH11R-NES-infiltrated leaves at 5 days post-infiltration (**Fig. 9B**; **Fig. S9D)**. Hence, the specific localization of RALPH11R at the nucleolus appears to be important for the induction of cell death, and its disruption prevents cell death in *N. benthamiana* leaves.

### Cell death induced by *Ep*RALPH11R is likely not due to HR activation

Plants generally respond to non-adapted pathogens by inducing ROS, pathogenesis-related genes, and localized cell death, a process known as the hypersensitive response (HR) (38). To determine whether cell death caused by RALPH11R overexpression is an HR response, we assessed hydrogen peroxide accumulation through DAB staining and expression of *PR* and HR-marker genes via RT-qPCR upon infiltration of GFP-RALPH11R, GFP-RALPH11F, GFP-*Fo*SRE1R (positive control), and empty vector (negative control) in *N. benthamiana* leaves. In contrast to the empty vector control, leaves infiltrated with GFP-*Fo*SRE1R, GFP-RALPH11F, and GFP-RALPH11R showed significantly higher DAB staining at 2 days post-infiltration, indicating that hydrogen peroxide accumulation is triggered by the full-length RALPH11 protein and its RNase domain (**Fig. 10A-B**). Similarly, the relative expression of *NbPR1* and *NbRBOHD* (gene encoding enzyme responsible for the generation of ROS) was found to be significantly upregulated in GFP-RALPH11F and GFP-RALPH11R-infiltrated leaves compared to the empty vector controls (**Fig. 10C)**. However, *NbHSR203*, a key marker gene for HR-associated cell death in plants (39), was significantly down-regulated in both RALPH11F and RALPH11R-infiltrated leaves compared to the empty vector controls (**Fig. 10C)**, indicating that the cell death observed upon RALPH11R overexpression is likely due to its RNase activity rather than HR induction.

**Figure 10.**
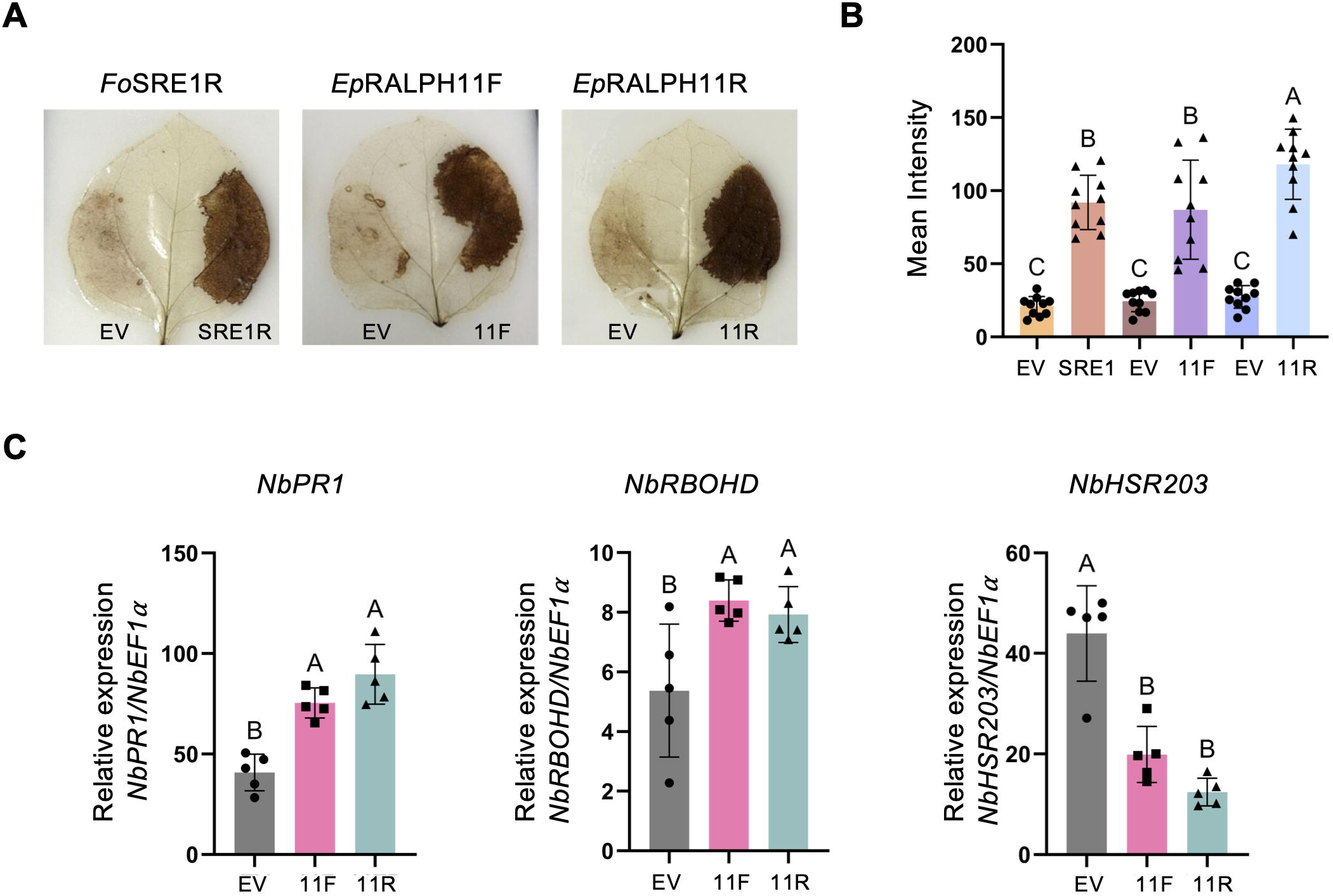
*Ep*RALPH11 full-length protein and its RNase domain induce defense responses, but not HR, in *N. benthamiana*. (A) Accumulation of Reactive oxygen species (ROS) 2 days post-infiltration of constructs expressing GFP-*Ep*RALPH11F, GFP-RALPH11R, *Fo*SRE1R (positive control), or GFP (EV, negative control) in *N. benthamiana* leaves. Leaves were stained with DAB (3,3 -Diaminobenzidine). (B) Quantitative analysis of DAB staining results in *N. benthamiana* leaves by ImageJ. Data represent mean ± SD (n=10). (C) Relative expression analysis of defense marker genes (*NbPR1, NbRBOHD*) and an HR marker gene (*NbHSR203*) 2 days post-infiltration of constructs expressing the indicated proteins in *N. benthamiana*. *NbEF1*α was used as a housekeeping gene for normalization. Data represent mean ± SD (n=5). Statistical significance was determined by an ordinary one–way ANOVA with Tukey’s multiple comparison test, where similar letters show non-significance.

### *Ep*RALPH11’s cell death-inducing ability may be suppressed by its N-terminal disordered region

Unlike its RNase domain, transient overexpression of the full-length RALPH11 protein did not induce cell death in *N. benthamiana* leaves. This was not surprising because CSEPs secreted by biotrophic PM pathogens typically suppress cell death when expressed in plants (40). BEC1054, a *Bgh* RALPH effector that lacks the catalytic residues required for RNase activity, has also not been reported to induce cell death in *N. benthamiana* (16). Unlike BEC1054, which is 118 amino acids long, RALPH11F is 225 aa long, consisting of a 104 aa long N-terminal structurally disordered region in addition to its 100 aa long RNase domain. To determine whether this structural feature affects the ability of this effector to induce cell death, we analyzed AF3 models of previously characterized RNase effectors from biotrophic and hemibiotrophic fungi (**Fig. 11A**). The comparative analysis showed that RNase effectors from hemibiotrophic fungi, including *Vd*RTX1 from *Verticillium dahliae*, *St*SRE1 from *Setosphaeria turcica,* and *Fg*12 from *Fusarium graminearum,* that induce cell death in *N. benthamiana*, are typically 132 to 138 amino acids long, consisting of a well-structured RNase domain (pLDDT Score>0.5) and a very small structurally disordered region (10-20 amino acids). However, like *Ep*RALPH11F, *Vd*RTX3, a 198 aa-long RNase-like effector secreted by *V. dahliae*, does not induce cell death despite possessing an RNase domain with conserved catalytic residues (35). *Vd*RTX3, like *Ep*RALPH11F, contains a structurally disordered region of the same length as that of its RNase domain, but at the C-terminal end (**Fig. 11A)**. Therefore, we speculate that the RNase activities of *Vd*RTX3 and *Ep*RALPH11F are suppressed or regulated by their extended intrinsically disordered regions. This regulation likely explains why *Ep*RALPH11F, and possibly *Vd*RTX3, fail to induce cell death in *N. benthamiana* despite harboring a catalytically active RNase domain (**Fig. 11B).**

**Figure 11.**
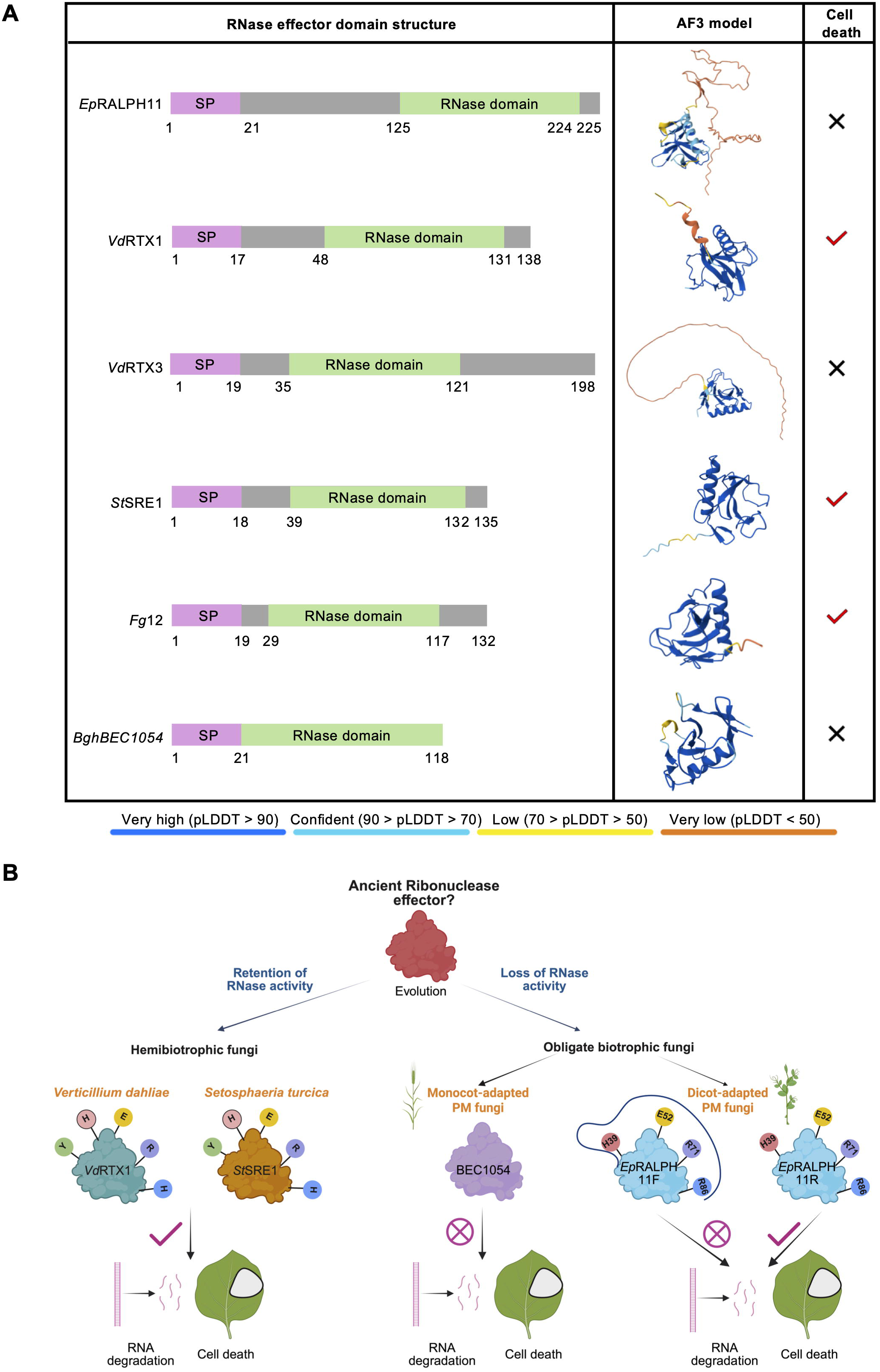
(A) AF3-predicted models of *Ep*RALPH11 and known RNase-like effectors from various hemibiotrophic and biotrophic fungal pathogens and their cell death-inducing ability in *N. benthamiana*. *Vd*RTX1 and *Vd*RTX3 effectors are from *Verticillium dahliae* (35), *St*SRE1 is from *Setosphaeria turcica* (36), *Fg*12 is from *Fusarium graminearum* (37), and BEC1054 is from *Blumeria graminis* f. sp. *Hordei* (16) (B) Proposed evolutionary model for the diversification of fungal RNase effectors. RNase effectors likely evolved from a common ancestral ribonuclease protein in both hemibiotrophic and obligate biotrophic fungi. In hemibiotrophs, RNase effectors retain the catalytic residues required for RNA degradation, enabling RNase activity and inducing host cell death, thereby promoting disease progression. In contrast, distinct evolutionary pressures associated with an obligate biotrophic lifestyle favored the attenuation or loss of RNase-mediated cytotoxicity. In cereal PMs, this was accompanied by the loss of catalytic residues essential for RNase activity, thereby rendering RALPHs catalytically inactive. In the pea PM *E. pisi*, RALPH11 retains key catalytic residues and an intrinsically active RNase domain capable of RNA degradation and cell death induction, like RNase effectors from hemibiotrophic fungi. However, its RNase activity is suppressed by the N-terminal structurally disordered region, which regulates the RNase domain and prevents host cell death, thereby enabling compatibility with the obligate biotrophic lifestyle. *Ep*RALPH11F, full-length; *Ep*RALPH11R, RNase domain

## Discussion

RALPH proteins with a catalytically inactive RNase domain constitute a major fraction of the effector repertoire of cereal PMs (9,13,41). RALPHs are also present in dicot-adapted PMs (15); however, unlike their counterparts in cereal PMs, they have remained largely uncharacterized. In the present study, we identified and compared the repertoires of RALPH CSEPs from *Erysiphe pisi* (*Ep*) and selected dicot-adapted PM fungi. We further performed a comprehensive structural and expression-based characterization of the *Ep*RALPHs and investigated the molecular and structural features associated with the function of an effector with a catalytically competent RNase domain.

Comparative analysis revealed that the RALPH CSEP family is undergoing expansion and diversification in the *Erysiphe* lineage. Firstly, the *Erysiphe* spp. contain a greater number of RALPHs, constituting ∼17-22% of the CSEPs compared to ∼1-12% in other dicot-adapted PM genomes (**Fig. 1A**; **Table S1**). Secondly, RALPHs with multiple RNase domains are found only in the *Erysiphe* genus, suggesting domain duplication events that may enhance the functional diversity of these proteins. Such multi-RNase-domain-containing RALPHs have not been reported previously in the cereal PMs. Thirdly, as shown for *E. pisi*, RALPHs that are physically clustered on the genome are also structurally related (**Fig. 1C**). Similar correlations between genetic and physical distance among RALPHs and other CSEP families have been reported in the *Blumeria* lineage (10), where gene duplication mediated by unequal crossing-over between flanking repeat sequences rich in specific transposable element (TE) families is proposed to have led to the expansion of these effector families (9,41). Although enrichment of TEs around CSEPs was not observed in the chromosome-level assembly of *E. necator* (42), genes encoding RNase-domain-containing proteins, including RALPH-like CSEPs, exhibit elevated duplication rates and are frequently organized in tandem clusters, a feature also observed in *E. pisi* (**Fig. 1C**). Given the recent or ongoing TE burst reported in *E. pisi* (43), TEs may have contributed to RALPH expansion in this lineage, although this remains to be demonstrated. The conserved intron position among tandemly arranged *Ep*RALPHs (**Fig. S6**) further supports their shared evolutionary origin.

Cereal PM *RALPHs* are typically expressed at very high levels in infected plants, predominantly in haustoria rather than in surface fungal spores and mycelia (9), suggesting that they perform key functions during fungal pathogenesis. *EpRALPHs* exhibit a similar expression pattern, with ∼75% of the analyzed *RALPHs* showing significantly higher expression in haustoria than in epiphytic mycelia (**Fig. 2A**; (25) Sharma et al. 2019). Many cereal PM *CSEPs*, including *RALPHs*, are expressed early during the infection process [e.g., (19)]. Likewise, almost all the analyzed *EpRALPHs*, except *EpRALPH18*, are induced during haustorium formation (between 12-24 hpi) (**Fig. 2B**), supporting a central role for these CSEPs in early pathogenesis.

Comparative AF3-based structural analysis indicates that *Ep*RALPHs contain well-structured RNase domain(s) flanked by intrinsically disordered regions at the N- and the C-termini (**Fig. 3**). Each *Ep*RALPH contains at least one and up to four RNase-like domains. Despite sharing a common RNase fold, these effectors exhibit significant sequence variation, suggesting functional diversification. Also, over the course of evolution, similarity with T1/F1 RNases has diminished, although the structural fold has been conserved (**Fig. 4**). The degree of similarity is mainly affected by local variations in the central β sheet, followed by differences in the peripheral β sheet (**Table S4**). The preservation of disulfide bonds contributes to the structural stability of these proteins, which may be crucial for maintaining the conserved structural architecture. Although the RNase domains show a similar degree of structural resemblance to T1 and F1 RNases, the similarity in the positions of disulfide bonds suggests that *Ep*RALPHs are evolutionarily more related to F1 than T1 RNases (**Table S5**). The majority of *Ep*RALPHs are predicted to lack RNase activity, as the canonical T1/F1 catalytic residues are absent from most family members. RALPHs 03 and 11 are notable exceptions, each retaining four of the six canonical T1/F1 catalytic residues, with RALPH11 predicted to exhibit a higher RNA-binding propensity than RALPH03 (**Fig. 5**; **Fig. S5**; **Table S6**). Nevertheless, most *Ep*RALPHs display a pronounced positive surface charge, distinct from that of T1/F1 RNases (**Fig. S7**), suggesting a potential nucleic acid-binding capability that may facilitate interactions with host RNA without requiring RNase catalytic activity, as observed with *Bgh* RALPHs (16). Further, the presence of multiple domains in some RALPHs may enhance the overall affinity of these effectors for their target nucleic acid or protein.

We selected RALPH11 for further functional characterization because it is the most highly expressed *Ep*RALPH and retains the core catalytic residues predicted to be required for RNase activity. In a previous study, we showed that full expression of *RALPH11*/*CSEP001* is required for the virulence of *E. pisi* on pea (25). Here, we show that transient overexpression of GFP-RALPH11 in a moderately susceptible Medicago genotype led to increased *E. pisi* hyphal formation compared with GFP-only controls, consistent with a role for RALPH11 in promoting host susceptibility to the pathogen (**Fig. 6A, B**). Transient expression assays revealed that RALPH11 localizes to the nucleolar rim of both host (*P. sativum*) and non-host (*N. benthamiana*) plants (**Fig. 6D**), suggesting that it may interact with rRNA and/or nucleolar proteins. The nucleolus is the primary site of rRNA synthesis and processing, where rRNAs are produced and assembled with ribosomal proteins to form the small and large ribosomal subunits (44). Nucleolar-targeting pathogen effectors can remodel the nucleolus by disrupting rRNA processing, ribosome biogenesis, and nucleolar protein distribution, thereby triggering nucleolar stress in plant cells (45,46). For example, a recent study showed that a nucleolar-localized *Phytophthora infestans* effector targets host ribosome biogenesis by disrupting pre-rRNA processing (46). We found that robust overexpression of RALPH11 in *N. benthamiana* alters the localization of the nucleolar protein Fibrillarin2, resulting in a fragmented nucleolar morphology (**Fig. 6E**). However, this nucleolar phenotype was not observed when RALPH11 was expressed in the host (*P. sativum*), possibly because of lower effector expression levels (**Fig. 6F**) or the presence of host-specific factors that mitigate its activity, thereby preventing the induction of detectable nucleolar stress. These findings suggest that RALPH11 targets host nucleolar functions to promote pathogenesis. Whether RALPH11 interacts with and interferes with the function(s) of nucleolar proteins, such as Fib2, remains to be investigated.

Despite sharing a common RNase scaffold with the T1/F1 RNases, cereal PM RALPHs lack the biochemical functions associated with these enzymes and are considered pseudo-RNases (12,13). In the present study, we demonstrate that the RNase domain of *Ep*RALPH11 (RALPH11R) degrades host RNA with specificity for single-stranded RNA (**Fig. 7**), suggesting that it retains RNase activity of the evolutionarily ancient RNase T1/F1 family. *Ep*RALPH11 may specifically target host ribosomal RNA, as evidenced by the reduction in 25S and 18S rRNA peaks following incubation of recombinant RALPH11R protein with pea total RNA (**Fig. 7**) and its nucleolar localization in plant cells (**Fig. 6**). As the RNase activity of full-length RALPH11 could not be assessed in vitro, we used transient expression in *N. benthamiana* as a proxy to compare the activities of the full-length (RALPH11F) and RNase-domain (RALPH11R) proteins. Such assays have previously been used to demonstrate the RNase activities of RNase-like effectors from hemibiotrophic fungal pathogens (35,36). We observed that expression of the RALPH11 RNase domain (11R), but not the full-length protein (11F), induced cell death in *N. benthamiana* leaves; however, this activity was abrogated when two of the four catalytic residues (H39 and R86) were mutated, suggesting that the cell death phenotype may result from RNA degradation (**Figs. 8 & 9**). In support of this, expression of *Ep*RALPH02 RNase domain, which lacks the conserved catalytic residues, did not induce cell death in *N. benthamiana*. Further, subcellular localization studies indicate that the cell death-inducing activity of RALPH11R depends on rRNA binding and/or cleavage, as the phenotype was associated with exclusive nucleolar localization of the RALPH proteins and was not observed when the proteins partitioned between the nucleolus and the nucleoplasm (RALPH11R^H39A^, 11R^R86A^, 11R^HERR-A^) or localized exclusively to the nucleoplasm (RALPH02R), potentially reducing its access to rRNA substrates. Together, these findings suggest that the RNase domain of RALPH11 is catalytically active and that residues H39 and R86 contribute to its rRNA-binding and/or RNA-degradation activities.

In addition to RNA degradation, activation of a hypersensitive response (HR) contributes to cell death in *N. benthamiana* leaves expressing RNase effectors from hemibiotrophic fungal pathogens (35,36). Expression of the *Verticillium dahliae* RNase effector*, Vd*RTX1, led to ROS accumulation and significant upregulation of marker genes associated with oxidative stress (*RBOHD*), defense (*PR1*), and HR (*HSR203*) in *N. benthamiana* leaves (35). Although only the RNase domain induced cell death in *N. benthamiana*, both RALPH11F and 11R induced ROS accumulation and the expression of *RBOHD* and *PR1* in *N. benthamiana* leaves; however, expression of the HR marker, *HSR203*, was downregulated compared with control leaves (**Fig. 10**). These findings are consistent with moderate immune activation, often observed upon heterologous expression of pathogen effectors in *N. benthamiana* (47) and suggest that cell death in RALPH11R-expressing leaves is due to RNA degradation rather than HR induction.

Catalytically active and cytotoxic ribonuclease effectors are generally secreted by hemibiotrophic phytopathogenic fungi, such as *Setosphaeria turcica, V. dahliae, Zymoseptoria tritici,* and *Colletotrichum orbiculare*, to either facilitate the biotrophic-to-necrotrophic transition or confer a competitive edge over other microbes in their ecological niche, thereby contributing to their overall fitness (35,36,48–50). Notably, their RNase activity was found to be indispensable for their cytotoxic function. In contrast, virulent effectors secreted by PM fungi are typically non-cytotoxic, consistent with their obligate biotrophic lifestyle, which depends on maintaining host cell viability. Accordingly, the well-characterized *Bg* lineage cereal PM RALPHs lack the catalytic residues conserved in the canonical T1/F1 RNases and are incapable of degrading RNA (12,13). The RNase activity-deficient *Bgh* RALPH BEC1054 was proposed to function instead by binding to and protecting host rRNA from plant ribosome-inactivating proteins, thereby preventing host cell death that would otherwise be induced (16). Here, we identified an *E. pisi* RALPH effector that harbors a catalytically active RNase domain with RNA-degrading capacity, yet the full-length protein is non-cytotoxic in plant cells, likely because its RNase activity is suppressed by an extended N-terminal intrinsically disordered region (**Fig. 11A**). From an evolutionary perspective, RALPHs are thought to have originated from an ancestral fungal ribonuclease, with RNase activity retained in some hemibiotrophic fungi but lost in biotrophic fungi through neofunctionalization (13). Building on this model, we propose that RALPH evolution in dicot-adapted PM fungi may have followed a distinct trajectory (**Fig. 11B**), potentially reflecting differences in the evolutionary pressures experienced by cereal and dicot-adapted PMs, which differ in their degree of host specialization (43). Rather than completely losing RNase activity through substitutions of catalytic residues, as in cereal PM RALPHs, some dicot PM RALPHs may have retained catalytically competent RNase domains while evolving intrinsically disordered regions that suppress or regulate their activity. Such regulation could preserve RNase-dependent functions while preventing excessive host cell damage, thereby supporting the obligate biotrophic lifestyle. Future studies examining the host protein interactors of *Ep*RALPH11 and the nucleolar processes it perturbs in the host plant will be important for understanding the biological significance of retaining a catalytically competent RNase domain in a dicot-adapted PM fungus.

## Materials and Methods

### In silico prediction of powdery mildew (PM) RALPH candidates

For the genome-wide identification of dicot-PM RALPHs, the protein databases of *Erysiphe pisi* (isolate Palampur-1), *E. necator* [isolate C-strain; (51)], *E. pulchra* [isolate Cflorida; (52)], *Oidium neolycopersici* [isolate UMSG2; (53)], *Golovinomyces cichoracearum* [isolate UMSG1 (53)], *G. orontii* (MGH1), and *Parauncinula polyspora* (54) were downloaded from MycoCosm (https://mycocosm.jgi.doe.gov/Erypi2/Erypi2.home.html). Protein sequences beginning with methionine and predicted to have a signal peptide (SP) with a D-Score>=0.5 were subjected to further analysis to identify candidate secreted proteins (CSPs), constituting the ‘secretome’, and candidate secreted effector proteins (CSEPs) as previously described (25). Briefly, TargetP (v2.0) (55) was used for the prediction of N-terminal presequences, and proteins predicted to have an ‘SP’ were retained. Prediction of transmembrane (TM) helices was performed by TMHMM (v2.0) (56), and proteins with no TM or TM within the SP were retained. Proteins predicted to have a GPI modification site using the Big-Pi Fungal predictor (57) were removed. EffectorP (v3.0; fungi and oomycetes) (58) was used to predict CSEPs from the respective secretomes. To identify candidate RALPHs, sequence- and structure-based analyses were performed. CSEPs predicted to have a ribonuclease domain/fold were identified using InterPro analysis (v8.30-90.0) (59). The structures of all CSEPs (minus SP) were predicted by ColabFold v1.5.5 (AlphaFold2 using MMseqs2) (60). For each protein, five models were generated, ranked from 001 to 005 based on the average pLDDT score and pTM score. We selected the model (ranked_001.pdb) with the highest average pLDDT score and pTM score for structural similarity analysis. Rupee was used for structural similarity comparisons against PDB chain databases (TOP_ALIGNED, FULL_LENGTH) downloaded on 16^th^ July 2022 (61). The overall bioinformatics pipeline is summarized in **Figure S1**. To predict the number of RNase domains per protein, RALPH sequences (minus SP) were modeled using AlphaFold 3 (AF3) with automatic seed 1 (27). Of the five models predicted for each RALPH, model_0 with the highest pLDDT and pTM scores was selected. The distribution patterns of introns and the scaffold locations of *EpRALPHs* were obtained from MycoCosm.

### Structural Phylogenetics

The AlphaFold 3-predicted CIF files (model_0 with the highest pLDDT and pTM values) for 126 dicot PM RALPHs were converted to PDB files using ChimeraX and submitted to the Foldtree server of Neurosnap (https://neurosnap.ai/service/Foldtree). The Foldtree output (.nwk file) was visualized and edited in iTOL7 (https://itol.embl.de/; (62)).

### Plant and fungal growth conditions and PM assays

*Pisum sativum* cv. AP-3 (pea), *Nicotiana benthamiana*, and *Medicago truncatula* (R108) seeds were grown in Conviron growth chambers at 22°C, 70% relative humidity, and a 16/8-h photoperiod with photosynthetically active radiation of 170 μmol m−2 s−1. *Fusarium oxysporum* f. sp. *ciceri* strain 7685 was grown on Potato Dextrose Agar Media (PDA) at 28°C for 5 days, followed by an additional 5 days in Potato Dextrose Broth in a shaker incubator set to 28°C and 100 rpm speed for optimum growth.

For the PM infection assays, leaves of 10–12-day-old pea plants and 3-week-old Medicago plants were inoculated with *Erysiphe pisi* isolate Palampur-1 (25) conidia from heavily infected AP-3 pea leaves [5–7 days post inoculation (dpi)] by the brushing and settling tower methods (63), respectively. To visualize fungal morphology, infected Medicago leaves were harvested 2 days post-inoculation and stained with trypan blue as described in (64). The stained leaves were mounted in 60% glycerol and observed with a 5× objective on a PALM MicroBeam microscope (Zeiss).

### Spatial and temporal RT-qPCR analyses

For the spatial expression analysis, heavily infected pea leaves (9 dpi) were painted with 5 % (w/v) cellulose acetate (Sigma, St. Louis, Missouri, USA) dissolved in 100% acetone (Sigma). Immediately after leaf drying, the fungal material trapped in cellulose acetate was gently peeled, frozen in liquid nitrogen, and used as the epiphytic sample. Pea leaves devoid of surface mycelium were frozen in liquid nitrogen and regarded as the haustorial sample. Total RNA was extracted from haustorial and epiphytic samples using the RNAiso Plus reagent (TaKaRa, Shiga, Japan), followed by DNase treatment with TURBO DNase (Thermo Fisher Scientific, Waltham, MA, USA). For the infection-dependent temporal expression analysis, *Ep*-infected pea leaves were harvested at 0, 12, 24, 48, 72, and 120 hpi and immediately frozen in liquid nitrogen. Total RNA was extracted from frozen samples using the Nucleospin RNA Plant kit (Macherey-Nagel, Dueren, Germany) with on-column DNase treatment to remove contaminating genomic DNA.

First-strand cDNA was synthesized from total RNA using PrimeScript 1^st^ strand cDNA synthesis kit (TaKaRa), and RT-qPCR was performed using TB green premix Ex Taq (Tli RNase H Plus) (TaKaRa) in a QuantStudio Flex 6 ABI system (Thermo Fisher Scientific). LinRegPCR was used to calculate mean PCR efficiencies per amplicon (65), and efficiency-corrected relative expression values were normalized to two reference genes, *Ep* β*-tubulin* 2 (*EpTUB2*, NCBI accession: X81961) and *Ep* 18S rRNA. The primer list is provided in **Table S7**.

### Structural analysis of *Ep*RALPH RNase domains

For each *Ep*RALPH, the amino acids forming well-structured α-helices and β-sheets in the AF3 model_0 were considered as the RNase domain. To determine the secondary structural elements (α-helices and β-sheets), the AF3 models were visualized in ChimeraX, and the command “dssp reports true” was selected. Depending on their position in the secondary structure, the β-sheets were categorized into central β-sheet (present in the central region and after the long α-helix) and peripheral β-sheet (present either at N-terminal or both N- and C-terminal). The number of β-strands within each β-sheet was recorded from the “dssp report true” outputs using the criteria of a minimum of 3 amino acids for each β-strand.

To assess similarity with canonical fungal F1 and T1 RNases, the *Ep*RALPH RNase domains (*Ep*RALPHR), modeled in AF3 (27), were structurally aligned with 2’-GMP-bound T1 (PDB:1RNT) and F1 (PDB:1FUT) RNases using FoldMason (66). The MSA LDDT scores, representing the degree of similarity, were recorded for each RALPH. The PDB files of the alignments were downloaded and visualized in ChimeraX version 1.10 (2025-06-26) (67). The electrostatic surface potential was generated in ChimeraX version 1.10.

To identify putative ligand binding residues, the *Ep*RALPHR protein sequences were submitted to the I-TASSER tool (http://zhanglab.ccmb.med.umich.edu/I-TASSER; (68). The C-Score, representing the reliability of the prediction, was recorded for each protein. To compare the positions of the ligand-binding residues in *Ep*RALPH11 with those in T1 RNase (PDB:1RNT), the I-TASSER model was energy minimized and sequence alignment was generated from structural alignments using TM-Align (69).

To identify the disulfide bond-forming cysteines, AF3 models of the *Ep*RALPH RNase domains were opened in ChimeraX, version 1.10. Cysteine residues were selected using the ‘Select-Residues-Cys’ option, visualized by choosing the ‘show atoms’ option, and disulfide bonds created using the command, “bond :cys@sg reasonable true”.

### Multiple sequence alignment

A multiple sequence alignment (MSA) using the amino acid sequences of *Ep*RALPH RNase domains was performed using Clustal Omega (70). The aligned sequences were graphically represented using the ESPript3 server (71).

### Plasmid construction and site-directed mutagenesis

The CDS (Coding DNA Sequence) minus SP for the full-length (F) and ribonuclease (R) domains of *Ep*RALPH11 (RALPH11F, RALPH11R) and *Ep*RALPH02 (RALPH02F, RALPH02R) were amplified from cDNA synthesized from total RNA isolated from *Ep*-infected pea leaves. The *Fo*SRE1 ribonuclease domain (*Fo*SRE1R) and *Ps*Fib2 CDS were amplified from cDNA prepared from RNA isolated from *Fusarium oxysporum* f. sp. *ciceri* strain 7685 and pea leaves, respectively. All amplified fragments were cloned into their respective expression/destination vectors using the Gateway BP Clonase II enzyme mix kit (Invitrogen, Cat No. 11789020) and the Gateway LR Clonase II enzyme mix kit (Invitrogen, Cat No. 11791020) according to the manufacturer’s instructions.

For localization and in-planta cell death assays, the destination vectors pSITE-2CA and pGWB654 were used to generate N-terminal GFP-tagged or C-terminal RFP-tagged constructs under the constitutive Cauliflower Mosaic Virus (CaMV) 35S promoter. For NES-tagging, the NES signal sequence (35) was fused to the C-terminus of RALPH11R and cloned into the pSITE-2CA vector. For protein expression and purification, the RALPH11R and RALPH02R sequences were cloned into pGEX-6P-1 (N-terminal GST tagging) by restriction digestion-based cloning. All primers used for cloning are listed in **Table S8**.

To generate the *Ep*RALPH11R mutant constructs, the predicted ligand-binding residues were mutated individually and in combination using a site-directed mutagenesis kit (Thermo Scientific, Cat No. F-541). The RALPH11R-pDONR207 plasmid was used as the initial template for the PCR reaction to generate the single (RALPH11R^H39A/E52A/R71A/R86A^) and quadruple (RALPH11R^HERR-A^) mutants using the primers listed in **Table S8**. PCR was performed using the following conditions: 25 cycles of 98°C for 10 sec, 71/72°C for 30 sec, and 72°C for 2 min. After PCR amplification, the parental template was digested with DpnI for 30 minutes, then transformed into *E. coli* DH5α competent cells. Positive mutants were screened through Sanger sequencing, and confirmed mutants were cloned into pSITE-2CA using the Gateway LR Clonase II enzyme mix kit (Invitrogen).

### Transient expression analysis and confocal microscopy

*Agrobacterium tumefaciens* EHA105 cells harboring the different plasmids were grown in LB medium at 28°C under appropriate antibiotic selection for 36 h. The cultures were centrifuged and resuspended in the infiltration buffer (10 mM MgCl_2,_ 10 mM MES at pH 5.6, and 150 mM Acetosyringone) to an OD of 0.8-1, then incubated at 28°C for 8-9 h to induce virulence before infiltration. The resuspended cultures were infiltrated into the abaxial leaf surface of 4-week-old *N. benthamiana*, 2-week-old *P. sativum*, or 3-week-old *M. truncatula* plants using a 1 ml needleless syringe. For the confocal assays where *Ps*Fib2 was used as the nucleolus marker, GFP-tagged constructs and *Ps*Fib2-RFP were co-infiltrated in a 1:1 ratio.

Imaging was performed on *N. benthamiana* leaves at 2 days post-infiltration and on *P. sativum* leaves at 2.5 days post-infiltration using a Leica TCS SP8 confocal microscope (Leica Microsystems, Germany). A two-channel sequence was used: TRITC for RFP (excitation peak: 561 nm, emission range: 570 nm-620 nm) and FITC for GFP (excitation peak: 488 nm, emission range: 500 nm-550 nm). Images were captured at 1024*1024 resolution, 200 speed, line average 4, and zoom factor 3. Images were analyzed and processed using LeicaX (LAS X, 3.5.7.23225). To quantify the nucleolar fragmentation phenotype, at least 100 nucleoli were recorded in GFP-RALPH11F- and empty vector (GFP)-infiltrated leaves across three independent experiments. Cell death symptoms were recorded at 5 days post-infiltration in leaves of *N. benthamiana* and *P. sativum*.

### Protein extraction and Western blotting

Agroinfiltrated leaf samples, harvested at 2 dpi (*N. benthamiana*) or 2.5 dpi (*P. sativum* and *M. truncatula*), were ground in liquid nitrogen using a mortar and pestle, resuspended in a protein extraction buffer containing 50 mM Tris-HCl, pH 7.5, 150 mM NaCl, 10% Glycerol, 10 mM MgCl_2,_ 1 mM DTT, 1 mM EDTA, and 1x Plant Protease Arrest Protease Inhibitor Cocktail (G-Biosciences, 786-332), and incubated on ice for 30 min. The samples were centrifuged at 14,000 rpm for 15 min at 4 °C, and the supernatant was mixed with 5X protein loading dye and boiled at 95 °C for 10 min. The samples were run on a 10% SDS-PAGE gel and transferred to a PVDF membrane using standard procedures. Blots were incubated in 0.5% skimmed milk (prepared in 1X TBST, pH 7.6) for 1 h, washed with 1X TBST solution twice, and incubated with 1.5:10,000 α-GFP antibody (Anti-GFP, N-terminal antibody produced in rabbit, G1544-100UG, Sigma-Aldrich), prepared in 1X TBST, at 4 °C overnight. Following primary antibody treatment, membranes were washed 6 times with 1X TBST (10 min each) and incubated with 1:10,000 Goat anti-Rabbit IgG (H & L) secondary antibody (HRP CONJUGATED, Agrisera, AS09 602), prepared in 1X TBST, for 1 h at room temperature. The blots were washed 6 times in 1X TBST solution and imaged using an ImageQuant LAS 4000 (GE Healthcare Life Sciences, USA).

### Detection of ROS and expression analysis of defense-related markers

For ROS detection, *N. benthamiana* and *P. sativum* leaves were stained with 1 mg/mL 3,3 -Diaminobenzidine (DAB) dye (Sigma-Aldrich, D8001-5G) at 2 (*N. benthamiana*) or 2.5 days (*P. sativum*) post-agroinfiltration. To dissolve DAB, the pH was adjusted to 3.5 with hydrochloric acid, then readjusted to pH 7 with 200 mM Na_2_HPO_4_. Leaves were incubated on a shaker for about 10-12 h in the dark at room temperature, decolorized using a solution of Ethanol: Glacial acetic acid: Glycerol (3:1:1) in a boiling water bath at about 95°C. Images were captured, and quantitative analysis of DAB staining intensity was performed through ImageJ (72).

For expression analysis of defense-related markers (*NbPR1, NbRBOHD, NbHSR203*), *N. benthamiana* leaves were harvested two days post-agroinfiltration, followed by RNA isolation, cDNA preparation, and RT-qPCR analysis as described previously. *NbEF1-*α was used as the endogenous control. Five biological replicates per experiment and two independent experiments were performed. Statistical significance was determined by ordinary one–way ANOVA with Tukey’s multiple comparison test, where similar letters show non-significance.

### Expression and purification of recombinant proteins

The RNase domain sequences of *Ep*RALPH11 and *Ep*RALPH02 were codon-optimized for the bacterial system and synthesized in the pGEX-6P-1 vector to generate N-terminal GST-tagged proteins. Expression of these proteins and GST alone was carried out in *Escherichia coli* BL21 cells. Cells were cultured in 5 mL Luria-Bertani (LB) media containing Carbenicillin (100 μg/mL) at 37 °C overnight. 50-70 μL of primary culture was inoculated into a 5 mL secondary culture containing carbenicillin (100 μg/ml), after which expression was induced with 0.5 mM IPTG (isopropyl β-D-1-thiogalactopyranoside) at 28 °C for 4 h. The pelleted cells were solubilized in the lysis buffer (50 mM Tris-HCl, pH 8.0, 500 mM NaCl, 10% glycerol, 1 mM DTT, and 1 mg/ml lysozyme) and incubated on ice for 30 min. The resuspended samples were sonicated (at 50% amplitude, with a cycle of 5 sec on and 10 sec off for 20 minutes) using a probe sonicator (Labman Scientific Instruments, India) and analyzed via SDS-PAGE for induction. After confirmation of induction, the GST-tagged proteins were purified from a 1 L culture using GST beads (G-Biosciences, Glutathione Resin, 786-310). Briefly, after induction, sonication, and centrifugation of cultures, the supernatants were passed through a column containing GST beads (Glutathione Resin, G-Biosciences, 786-310). The beads were washed first with a low salt washing buffer (50 mM Tris-HCl, pH 8.0, 300 mM NaCl, 10% glycerol, 1 mM DTT) three times, followed by the high salt washing buffer (50 mM Tris-HCl, pH 8.0, 500mM NaCl, 10% glycerol, 1 mM DTT) three times. Finally, the protein was eluted from the beads using the elution buffer (50 mM Tris-HCl, pH 8.0, 300 mM NaCl, 10% glycerol, 1 mM DTT, 20 mM GSH, pH 8.0). The eluted samples were pooled and dialyzed overnight at 4 °C with constant stirring in dialysis buffer (50 mM Tris-HCl, pH 8.0, 300 mM NaCl, 10% glycerol). The dialyzed samples were concentrated using a concentrator (Amicon Ultra Centrifugal Filter, 10 kDa MWCO, Merck, UFC5010). The protein concentration was estimated using gel-based quantification using BSA as a standard. To check the purity of the protein, 1 µg of each purified protein (RALPH11R, RALPH02R, and GST) was loaded and run on a 10% SDS-PAGE gel, stained with Coomassie Brilliant Blue R-250 (Bio-Rad), destained, and visualized on a Gel Doc^TM^ EZ Imager (Bio-Rad).

### *In vitro* RNase activity assays

To test the ribonuclease activity of *Ep*RALPH11R, an *in vitro* RNase assay was performed as described in (12) with slight modifications. Total RNA was isolated from the leaves of three-week-old pea plants using the Macherey-Nagel (MN) Nucleospin RNA Plant kit (Germany) with on-column DNase digestion to remove genomic DNA. Purified GST-RALPH11R protein was incubated with pea leaf total RNA. The reaction mixture consisted of 1.8 µg RNA and 25 µM protein in a buffer containing 15 mM Tris-HCl (pH 8.0), 15 mM NaCl, 50 mM KCl, 2.5 mM EDTA, and nuclease-free water in 30 µl total volume. The reaction mixture was incubated at 25° C for 1 h. For analysis using the TapeStation System 4150 (Agilent, CA, USA), 1 µl of the reaction mixture was used. Purified RALPH02R, GST, and nuclease-free water (no protein) were used as the negative controls, while RNase T1 (Thermo Scientific, EN0541) was used as the positive control.

To confirm the specificity of *Ep*RALPH11R for single-stranded RNA, the above *in vitro* RNase assay was performed with GFP double-stranded RNA (dsRNA) as the substrate. GFP dsRNA was synthesized using a T7 in vitro transcription kit. Briefly, a 250 bp fragment of eGFP (pSITE-2CA vector) was PCR amplified using GFP-specific primers appended with T7 promoter sequences (T7-GFPFWD: <u>TAATACGACTCACTATAGGG</u>ATGCCCGAAGGCTACGTCCAG; T7-GFPREV: <u>TAATACGACTCACTATAGGG</u>TGTTGTGGCGGATCTTGAAG). Amplified products were eluted using FavorPrep™ GEL/PCR Purification Kit (Favorgen Biotech Corp., Taiwan). One microgram of PCR-eluted product was used as the template for dsRNA synthesis using the MEGAscript™ T7 Transcription Kit (Catalog no: AM1334; Invitrogen) according to the manufacturer’s protocol. Purified *Ep*RALPH11R (25 µM) was incubated with 1.8 µg GFP dsRNA or pea total RNA in the same reaction buffer mentioned above in 30 µl total volume and incubated at 25 ° C for 1 h. On completion of the reaction, 1 µL of the mixture was analyzed in an Agilent TapeStation System 4150. RNase A (Thermo Fisher Scientific, EN0531) and RNase T1 were used as positive controls, and GST and nuclease-free water were used as negative controls.

### Statistical analysis

Statistical analyses were performed using GraphPad Prism v11.0 (GraphPad Software, La Jolla, CA, USA).

## Supporting information

Workbook S1

## Data Availability

All relevant data are within the manuscript and its Supporting Information files.

## Declaration of generative AI and AI-assisted technologies in the manuscript preparation process

During the preparation of this manuscript, the authors used ChatGPT and Grammarly for language polishing, grammar checking, and readability improvement. After using these tools, the authors reviewed and edited the content as needed and take full responsibility for the article’s content.

## Acknowledgments

This work was supported by UGC-PhD Fellowships to Debashish Sahu, Smritilekha Mukherjee, and Poonam Ray, DBT-PhD fellowship to Puja Ghosh, and RCB Ramachandran DBT fellowship to Gagan Gupta. The sequence data used for the analysis were produced by the US Department of Energy Joint Genome Institute CSP Project 1657, led by Mary Wildermuth at UC Berkeley and in collaboration with the user community. We thank Dr. Praveen Verma at JNU for providing the Fusarium strain and Tanishka Agrawal for technical assistance with optimizing the RNase assays.

## Author contributions

**Conceptualization:** Deepti Jain, Divya Chandran.

**Data curation:** Debashish Sahu, Divya Chandran.

**Formal analysis:** Debashish Sahu, Puja Ghosh, Smritilekha Mukherjee, Vineet Kumar, Gunjan Sharma, Deepti Jain, Divya Chandran.

**Funding acquisition:** Deepti Jain, Divya Chandran.

**Investigation:** Debashish Sahu, Puja Ghosh, Smritilekha Mukherjee, Vineet Kumar, Gunjan Sharma, Megha Gupta, Gagan Gupta, Poonam Ray, Kusum, Jaya Sharma.

**Methodology:** Debashish Sahu, Puja Ghosh, Smritilekha Mukherjee, Deepti Jain, Divya Chandran.

**Project administration:** Debashish Sahu, Deepti Jain, Divya Chandran.

**Supervision:** Deepti Jain, Divya Chandran.

**Validation:** Debashish Sahu, Puja Ghosh, Deepti Jain, Divya Chandran.

**Visualization:** Debashish Sahu, Puja Ghosh, Smritilekha Mukherjee, Gunjan Sharma, Deepti Jain, Divya Chandran.

**Writing– original draft:** Debashish Sahu, Puja Ghosh, Smritilekha Mukherjee, Gunjan Sharma, Deepti Jain, Divya Chandran.

**Writing– review & editing:** Debashish Sahu, Puja Ghosh, Smritilekha Mukherjee, Vineet Kumar, Gunjan Sharma, Megha Gupta, Gagan Gupta, Poonam Ray, Kusum, Jaya Sharma, Deepti Jain, Divya Chandran.

## Supplementary Material

**Workbook S1.** List of RALPH CSEPs identified from different dicot-adapted PM species

**Table S1.** RALPH to CSEP ratios in different dicot-infecting PM species

**Table S2.** BLASTp analysis of RALPH-like CSEPs identified from the *Ep* transcriptome versus the genome

**Table S3**. Table showing the distribution of the number of RNase domains in *Ep*RALPHs

**Table S4**. Table showing the number of β-strands in *Ep*RALPHs compared to T1 and F1 RNases

**Table S5**. Distribution of disulfide bonds in *Ep*RALPHs compared to T1 and F1 RNases

**Table S6**. I-TASSER predictions for ligand and ligand-binding residues for *Ep*RALPH RNase domains with associated C-scores.

**Table S7**. List of RT-qPCR primers

**Table S8**. List of primers used in cloning and site-directed mutagenesis

**Figure S1.** *In silico* pipeline for identifying PM RALPH effector candidates

**Figure S2.** AlphaFold 3 (AF3) models of single and multi-RNase domain *Ep*RALPH full-length proteins

**Figure S3.** CDS of multi-RNase domain *EpRALPHs* amplified from *Ep* cDNA versus that predicted in MycoCosm

**Figure S4**. Exon-intron structures of predicted versus PCR amplified multi-RNase domain *EpRALPHs*

**Figure S5**. Multiple sequence alignment of RNase domains of *Ep*RALPHs along with T1 and F1 RNases, highlighting conservation of cysteine (asterisks) and catalytic residues (triangles).

**Figure S6**. Exon-intron structures of *Ep*RALPHs and multiple sequence alignment of single intron *Ep*RALPH RNase domains showing conservation of the intron position **Figure S7**. Surface charge potential of *Ep*RALPH RNase domains

**Figure S8**. (A) SDS-PAGE showing recombinant GST-*Ep*RALPH11R, GST-*Ep*RALPH02R and GST protein (B) TapeStation electropherograms for RNase activity reactions of different proteins with ssRNA (pea RNA) as substrate

**Figure S9.** Whole leaf images showing cell death symptoms after agroinfiltration of wildtype or mutant constructs of *Ep*RALPHs in *Nicotiana benthamiana* and *Pisum sativum* leaves.

## Notes

**Funding**: This study was supported by Anusandhan National Research Foundation-Science and Engineering Board Core Research Grant CRG/2020/003413 to DC and DJ. The funders had no role in the design of the study, in the collection, analyses, or interpretation of data, in the writing of the manuscript, or in the decision to publish the results.

### Competing Interest Statement

The authors have declared no competing interest.

### Summary of Updates

Supplemental Files attached; author order updated

