## Supplementary Information for "An extended N-terminus restrains the plant cell death-inducing ability of the catalytically competent ribonuclease domain in a pea powdery mildew RALPH effector"

**Table S1**. RALPH to CSEP ratios in different dicot-infecting PM species

| PM species | % RALPHs to total CSEPs |
|---|---|
| *Ep* | 18.3 |
| *En* | 21.7 |
| *Epul* | 17.3 |
| *On* | 7.8 |
| *Gc* | 11.3 |
| *Go* | 12.4 |
| *Pp* | 1.4 |

**Table S2.** BLASTp results of Ribonuclease-like EpCSEPs identified in the *Ep* haustorial transcriptome (Sharma, Aminedi et al. 2019) versus the *Ep* genome

| ***Ep*CSEP ID** | **Query length** | **Subject ID** | ***Ep*RALPH ID** | **% identity** | **Subject coverage %** | **length** | **mismatch** | **gapopen** | **qstart** | **qend** | **sstart** | **send** | **evalue** | **bitscore** |
|---|---|---|---|---|---|---|---|---|---|---|---|---|---|---|
| CSEP001 | 225 | Erypi2\|2358942 | RALPH11 | 100.00% | 100.00% | 225 | 0 | 0 | 1 | 225 | 1 | 225 | 1.18E-164 | 1234 |
| CSEP002 | 138 | Erypi2\|2358943 | | 99.18% | 66.30% | 122 | 1 | 0 | 1 | 122 | 1 | 122 | 6.90E-84 | 654 |
| CSEP009 | 176 | Erypi2\|858556 | | 100.00% | 71.40% | 135 | 0 | 0 | 1 | 135 | 1 | 135 | 2.89E-90 | 700 |
| CSEP009 | 176 | Erypi2\|2358953 | RALPH13 | 43.20% | 92.90% | 169 | 96 | 0 | 8 | 176 | 13 | 181 | 1.02E-43 | 365 |
| CSEP012 | 256 | Erypi2\|2710487 | RALPH23 | 98.28% | 60.50% | 116 | 2 | 0 | 125 | 240 | 108 | 223 | 2.58E-76 | 602 |
| CSEP012 | 256 | Erypi2\|2710487 | RALPH23 | 100.00% | 60.50% | 97 | 0 | 0 | 1 | 97 | 1 | 97 | 3.23E-65 | 522 |
| CSEP020 | 181 | Erypi2\|2652532 | RALPH19 | 100.00% | 100.00% | 181 | 0 | 0 | 1 | 181 | 1 | 181 | 1.18E-128 | 962 |
| CSEP023 | 209 | Erypi2\|1848142 | RALPH09 | 99.52% | 100.00% | 209 | 1 | 0 | 1 | 209 | 1 | 209 | 1.01E-143 | 1092 |
| CSEP027 | 217 | Erypi2\|2514510 | RALPH28 | 27.78% | 34.60% | 72 | 52 | 0 | 103 | 174 | 104 | 175 | 5.65E-07 | 101 |
| CSEP036 | 198 | Erypi2\|739599 | RALPH04 | 100.00% | 100.00% | 198 | 0 | 0 | 1 | 198 | 1 | 198 | 9.18E-139 | 1056 |
| CSEP036 | 198 | Erypi2\|1848134 | RALPH08 | 96.46% | 100.00% | 198 | 7 | 0 | 1 | 198 | 1 | 198 | 2.63E-131 | 1002 |
| CSEP039 | 182 | Erypi2\|2358953 | RALPH13 | 98.35% | 100.00% | 182 | 3 | 0 | 1 | 182 | 1 | 182 | 3.38E-121 | 933 |
| CSEP045 | 225 | Erypi2\|738751 | RALPH03 | 82.67% | 100.00% | 225 | 39 | 0 | 1 | 225 | 1 | 225 | 2.02E-134 | 1019 |
| CSEP045 | 225 | Erypi2\|2516098 | RALPH17 | 82.67% | 100.00% | 225 | 39 | 0 | 1 | 225 | 1 | 225 | 2.02E-134 | 1019 |
| CSEP066 | 110 | Erypi2\|2659066 | | 55.17% | 20.80% | 87 | 39 | 0 | 20 | 106 | 1 | 87 | 7.12E-30 | 260 |
| CSEP068 | 225 | Erypi2\|2358942 | RALPH11 | 82.22% | 100.00% | 225 | 40 | 0 | 1 | 225 | 1 | 225 | 6.05E-133 | 1008 |
| CSEP069 | 145 | Erypi2\|2660137 | RALPH20 | 99.19% | 67.80% | 124 | 1 | 0 | 1 | 124 | 1 | 124 | 4.20E-84 | 653 |
| CSEP102 | 148 | Erypi2\|2391848 | | 100.00% | 51.90% | 41 | 0 | 0 | 108 | 148 | 39 | 79 | 7.08E-23 | 214 |

| **Table S3**. Table showing the distribution of the number of RNase domains in *Ep*RALPHs | | | | |
|---|---|---|---|---|
| No. of RNase domains | One RNase domain | Two RNase domains | Three RNase domains | Four RNase domains |
| *Ep*RALPH ID | 1,2,3/17,4,5,8,9,11,12,13,14,15,16,19,20,24,25,26,27,28,29,30 | 18,21 | 22,23 | 6,7,10 |

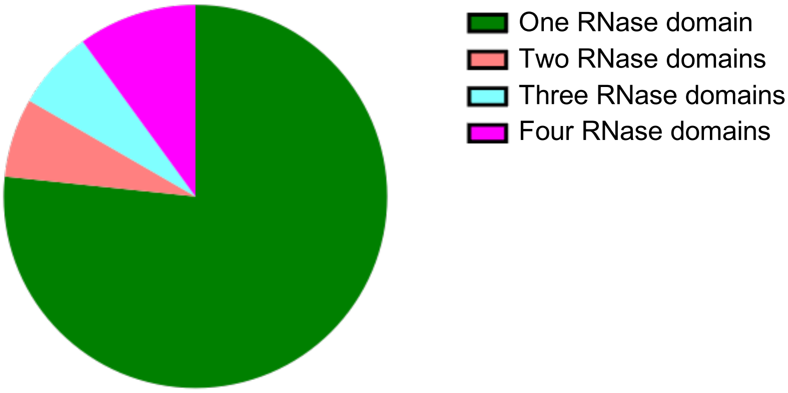

**Table S4**. The number of β strands present in *Ep*RALPHs compared to T1 and F1 RNases. Here, N and C denote the N-terminal and C-terminal regions, respectively.

| RALPH ID | Central β sheet | Peripheral β sheet |
|---|---|---|
| RALPH01 | 3 | 2 (2N) |
| RALPH02 | 4 | 2 (2N) |
| RALPH03 | 5 | 3 (2N+1C) |
| RALPH04 | 5 | 3 (2N+1C) |
| RALPH05 | 3 | 0 |
| RALPH06A | 4 | 3 (2N+1C) |
| RALPH06B | 4 | 2 (2N) |
| RALPH06C | 5 | 2 (2N) |
| RALPH06D | 5 | 0 |
| RALPH07A | 4 | 2 (2N) |
| RALPH07B | 5 | 2 (2N) |
| RALPH07C | 5 | 0 |
| RALPH07D | 6 | 0 |
| RALPH08 | 5 | 3 (2N+1C) |
| RALPH09 | 5 | 0 |
| RALPH10A | 4 | 2 (2N) |
| RALPH10B | 4 | 0 |
| RALPH10C | 5 | 3 (2N+1C) |
| RALPH10D | 6 | 0 |
| RALPH11 | 5 | 3 (2N+1C) |
| RALPH12 | 3 | 2 (2N) |
| RALPH13 | 5 | 0 |
| RALPH15 | 5 | 3 (2N+1C) |
| RALPH16 | 4 | 2 (2N) |
| RALPH18A | 4 | 2 |
| RALPH18B | 5 | 3 (2N+1C) |
| RALPH19 | 6 | 2 (2N) |
| RALPH20 | 5 | 3 (2N+1C) |
| RALPH21A | 5 | 2 (2N) |
| RALPH21B | 4 | 3 (2N+1C) |
| RALPH22A | 4 | 2 (2N) |
| RALPH22B | 4 | 2 (2N) |
| RALPH22C | 3 | 2 (2N) |
| RALPH23A | 4 | 2 (2N) |
| RALPH23B | 4 | 2 (2N) |
| RALPH23C | 3 | 2 (2N) |
| RALPH24 | 7 | 0 |
| RALPH25 | 5 | 0 |
| RALPH26 | 5 | 3 (2N+1C) |
| RALPH27 | 3 | 0 |
| RALPH28 | 4 | 2 (2N) |
| RALPH29 | 4 | 3 (2N+1C) |
| RALPH30 | 4 | 2 (2N) |
| T1 RNase | 5 | 2 (2N) |
| F1 RNase | 5 | 2 (2N) |

**Table S5**. Distribution of disulfide bonds in *Ep*RALPHs compared to T1 and F1 RNases

| RALPH ID | T1/F1 RNase | F1 RNase | T1 RNase | New |
|---|---|---|---|---|
| RALPH01 | 1 | - | - | - |
| RALPH02 | 1 | 1 | - | 1 |
| RALPH03 | 1 | 1 | - | - |
| RALPH04 | 1 | 1 | - | - |
| RALPH05 | - | - | - | - |
| RALPH06A | 1 | - | - | - |
| RALPH06B | 1 | - | - | - |
| RALPH06C | 1 | - | - | - |
| RALPH06D | 1 | 1 | - | - |
| RALPH07A | 1 | - | - | - |
| RALPH07B | 1 | - | - | - |
| RALPH07C | - | - | - | - |
| RALPH07D | 1 | 1 | - | - |
| RALPH08 | 1 | 1 | - | - |
| RALPH09 | 1 | 1 | - | - |
| RALPH10A | 1 | - | - | - |
| RALPH10B | 1 | - | - | - |
| RALPH10C | 1 | - | - | - |
| RALPH10D | 1 | 1 | - | - |
| RALPH11 | 1 | 1 | - | - |
| RALPH12 | - | - | - | - |
| RALPH13 | 1 | 1 | - | - |
| RALPH15 | 1 | 1 | - | - |
| RALPH16 | 1 | 1 | - | - |
| RALPH18A | 1 | - | - | - |
| RALPH18B | 1 | 1 | - | - |
| RALPH19 | 1 | 1 | - | 1 |
| RALPH20 | 1 | 1 | - | - |
| RALPH21A | 1 | - | - | - |
| RALPH21B | 1 | 1 | - | - |
| RALPH22A | 1 | - | - | - |
| RALPH22B | 1 | - | - | - |
| RALPH22C | - | - | - | - |
| RALPH23A | 1 | - | - | - |
| RALPH23B | 1 | - | - | - |
| RALPH23C | - | - | - | - |
| RALPH24 | 1 | 1 | - | - |
| RALPH25 | 1 | 1 | - | - |
| RALPH26 | 1 | 1 | - | 1 |
| RALPH27 | - | 1 | - | - |
| RALPH28 | 1 | 1 | - | 1 |
| RALPH29 | 1 | - | - | - |
| RALPH30 | 1 | 1 | - | 1 |
| T1 RNase | 1 | - | 1 | - |
| F1 RNase | 1 | 1 | - | - |

**Table S6**. I-TASSER predictions for ligand and ligand-binding residues for *Ep*RALPH RNase domains with associated C-scores. Known catalytic residues in T1/F1 RNases and corresponding putative residues in *Ep*RALPHs are highlighted in bold. Here, 2'-GMP is abbreviated as 2GP.

| **RALPH ID** | **Ligand name** | **C-score** | **Ligand binding-site residues** |
|---|---|---|---|
| T1 RNase | 2GP | 0.99 | **Y38**,**H40**,41,42,43,44,45,46,**E58**,**R77**,**H92**,98,**F100** |
| F1 RNase | 2GP | 0.99 | **Y38**,**H40**,41,42,43,44,45,46,**E58**,**R76**,**H91**,97,98,**F99** |
| RALPH01 | 2AM | 0.73 | 35,37,39,49,68,69,**R71**,86,92,94,**F98** |
| RALPH02 | 3GP | 0.39 | 38,40,43,44,45,46,47,50,61,84,99,105,106,107 |
| RALPH03 | GPC | 0.66 | 35,37,38,**H39**,40,51,**E52**,67,68,69,**R71**,**R86**,89,90,92 |
| RALPH04 | 4GSPA00 | 0.18 | 36,39,41,**R86**,92,93,94 |
| RALPH05 | 2GP | 0.40 | **Y27,**29,30,31,32,34,36,37,56,76,92 |
| RALPH06A | 2GP | 0.48 | **Y33**,35,36,37,38,39,40,41,52,73,89,93,95 |
| RALPH06B | 2GP | 0.47 | **Y31**,33,34,35,36,37,48,69,**R84**,88,89,**F90** |
| RALPH06C | 2AM | 0.31 | 35,37,50,65,66,67,**R69**,84,86,89 |
| RALPH06D | 2GP | 0.41 | **Y32**,34,35,36,37,38,39,49,**R72**,87,93,94 |
| RALPH07A | 2GP | 0.48 | **Y31**,33,34,35,36,37,38,50,51,**R87**,91,92,94 |
| RALPH07B | 2GP | 0.58 | **Y31**,33,34,35,36,37,38,48,73,**R88,**92,93,94 |
| RALPH07C | 2GP | 0.39 | 19,21,22,23,25,26,34,**R53**,68,71,73 |
| RALPH07D | 1GSPA00 | 0.13 | **Y32**,36,38,79,**R88**,115,116,117 |
| RALPH08R | VO4 | 0.26 | 37,39,52,**R71**,**R86**,92,94 |
| RALPH09R | 2GP | 0.17 | 33,35,36,37,38,40,41,**E50,R96,R111**,116,117 |
| RALPH10A | 2GP | 0.50 | **Y31,**33,34,35,36,37,38,50,**R71**,87,91,92,93 |
| RALPH10B | 2AM | 0.47 | **Y30,**32,47,65,68,83,87,89 |
| RALPH10C | 2AM | 0.35 | 36,38,51,66,67,68,**R70,**85,88,90 |
| RALPH10D | 2GP | 0.68 | **Y32**,34,35,36,37,38,39,49,**R66,**81,88,89,90 |
| RALPH11 | 2GP | 0.95 | 35,37,38,**H39**,40,41,**E52,R71,R86**,90,91,92 |
| RALPH12 | 2AM | 0.19 | 34,36,**E51**,72 |
| RALPH13 | 2GP | 0.87 | 33,35,36,37,38,39,**E50,R69,R84**,88,90 |
| RALPH15 | 2GP | 0.25 | 44,**Y45,**46,**E59,R82**,97,101,103 |
| RALPH16 | 2GP | 0.38 | 39,40,41,47,48,65,89,104,108,109,**F110** |
| RALPH18A | 2GP | 0.32 | 32,35,37,50,64,65,66,**R69**,84,87,89 |
| RALPH18B | 2GP | 0.59 | **Y34**,36,37,38,39,40,41,52,**R71**,86,90,91,92 |
| RALPH19 | 2GP | 0.40 | **Y52**,54,55,56,57,58,59,60,78,**R98**,113,119,120 |
| RALPH20 | 2GP | 0.41 | 39,**Y41,**42,43,44,45,46,47,59,76,91,97,98 |
| RALPH21A | 2GP | 0.31 | 42,43,44,45,46,56,**R76**,91,96 |
| RALPH21B | 2GP | 0.85 | 35,37,38,39,40,41,43,45,57,76,91,100,102 |
| RALPH22A | 2GP | 0.52 | **Y31**,33,34,35,36,38,39,50,**R71**,87,91,92,93 |
| RALPH22B | 2GP | 0.36 | 31,33,34,35,37,38,48,69,84,88,89,90 |
| RALPH22C | 2ql2A | 0.15 | 17,18,21 |
| RALPH23A | 2GP | 0.58 | **Y37,**39,40,41,42,43,44,45,56,**R77**,93,97,98,99 |
| RALPH23B | 2GP | 0.53 | 31,33,34,35,36,37,38,48,69,84,88,89,90 |
| RALPH23C | 2GP | 0.12 | 35,36,**Y37,**38 |
| RALPH24 | 2GP | 0.63 | **Y50**,52,53,54,55,56,57,58,69, 90,104,108,110 |
| RALPH25 | 2GP | 0.53 | 33,35,36,37,38,39,40,**E50,R61,R76**,80,82 |
| RALPH26 | 2GP | 0.61 | **Y40**,42,43,44,45,46,47,48,61,82,97,103,110,111 |
| RALPH27 | 2AM | 0.23 | 47,49,51,71,86,87,89,104,114,115,118 |
| RALPH28 | 1BVIA00 | 0.20 | 36,37,38,40,41,45,46,49,**R87**,110,112 |
| RALPH29 | 1LOVA00 | 0.24 | 39,42,43,44,45,77,97,100,101,102,103,104,106 |
| RALPH30 | 2GP | 0.22 | 41,53,54,55,86,105,110 |

| **Table S7.** List of RT-qPCR primers used in the study. | | | |
|---|---|---|---|
| Gene Name | Protein ID | Forward Primer (5'-3') | Reverse Primer (5'-3') |
| *EpRALPH01* | 55905 | AAGCTCGTTGCGGCTCTTAT | ACCATCAGTTCCAAAGGGCT |
| *EpRALPH02* | 517564 | GTGAGCCAATGAAGGTTGACG | TCGTCCTACCCTTGGGTTCT |
| *EpRALPH03*<br>*EpRALPH17* | 738751<br>2516098 | ATGGACGGTTCTACACCAGC | TGTGTCGGACTACTGCTCCT |
| *EpRALPH04* | 739599 | GCAGCCTGTCCTCGCATATC | CAAAACCTGCCGTTACGCTG |
| *EpRALPH05* | 824921 | TAGAGCCAACAGATCCAGCAG | ATACAAGGCAGCGCCACAT |
| *EpRALPH06* | 1813428 | TGGTAGGTTATGAGTGCGGC | GCCTCGGTATTGGCTAGGAT |
| *EpRALPH07* | 1847045 | AGGAAGGTTATGAGTGCGGC | TCACCGAAAAGTTTGCCTCG |
| *EpRALPH08* | 1848134 | TCAAGATTGACCCAGCGGAC | CCACATCGAAGGCCGTATCT |
| *EpRALPH09* | 1848142 | CTCGCTCTATACCGCAAGCA | GCTCGGGAAGTCCCAATACT |
| *EpRALPH10* | 2358292 | CCCATCACCTACGGTTCCAG | GGGCCAAGGCAACACTTCTA |
| *EpRALPH11* | 2358942 | TTGACAACGACGGGCCTTAC | ACCCTTGTGTCGGACTACTG |
| *EpRALPH12* | 2358946 | CCTTTGACAAGTACGGGCCT | CGGGGCAGTAGGGACTTTTC |
| *EpRALPH13* | 2358953 | AGAGTGGCCAATTTCGCGT | TCACGGCACCAACCACTTTA |
| *EpRALPH14* | 2361005 | GCTTTCCCACTACTCGCCAA | ATACCCGCAACTTCCCCATC |
| *EpRALPH15* | 2362051 | GGCTCAAAGCATCAGCGAAC | TGGACCAAGAAATCTGGCGTA |
| *EpRALPH16* | 2430685 | TATTTGCGGGCCAGTGTAGG | TAGGCCGTATAGGGAGGTGG |
| *EpRALPH18* | 2586088 | TCTCCTGTATTGCCCGAAGC | GGCGGACCATTGTATGCTTTC |
| *EpRALPH19* | 2652532 | CACTATTGGAGTTTGGTCGCA | ACATGGCTCAGTAGAGCGTC |
| *EpRALPH20* | 2660137 | TGCTGCTGAGACTGCTTGTA | ATTGCCGGGGATTTCGAATG |
| *EpRALPH21* | 2672905 | ATTTTCCGAGGGTGATCGCA | GGTAGCCTTGCAGCCATTTC |
| *EpRALPH22* | 2710286 | ACCGCATTTCGTGTTGTCTT | TGGCCCTATGTTCGTACTGC |
| *EpRALPH23* | 2710487 | TCGACCGCATACCAAAACCA | TATGTTCGTACTGCCTCGCC |
| *EpRALPH24* | 2851786 | GCTTTCCCACTACTCGCCAA | ATACCCGCAACTTCCCCATC |
| *EpRALPH25* | 2863788 | TGATCCGGGGGAAGAACAGT | AGCTAATTTCTTGGCTTCTCTCTCA |
| *EpRALPH26* | 652229 | CGACGTTTCTGGCGAAGAAT | GCACGCCGAAGATTCACATT |
| *EpRALPH27* | 2375095 | ACTACGGTTCAAGTACGGGC | ACCCTTCCTGTTTGTCCTTCG |
| *EpRALPH28* | 2514510 | AAAGGGGCTGCAATGTTTGG | TAGCCCGTTCGAGGCAAATC |
| *EpRALPH29* | 2530120 | GTACCGAGCCAAGGCGAAAA | CTCCATGCAAATCCTGGTGC |
| *EpRALPH30* | 2601571 | AAGGGCCTCTGTGTGGTAAAG | GAGTGCCGTGGCTTTATTCC |
| *EpTUB2* | | ATCTGCCGTCATGTCAGGTG | CAGTGACAGCCCGGAATGAG |
| *Ep18S rRNA* | | TCCAGCTGCCTTTGTGTGG | GATGAGGGTTGTTCTGGCAAGC |
| *NbRBOHD* | | TTTCTCTGAGGTTTGCCAGCCACCA | GCCTTCATGTTGTTGACAATGTCTTT |
| *NbPR1* | | CCGCCTTCCCTCAACTCAAC | GCACAACCAAGACGTACTGAG |
| *NbHSR203* | | GGCAGTGGAGGAGCTTAAAT | GCTATGTCCCACTCCATTGTTA |
| *NbEF1*$\alpha$ | | TGAGTTCGAGGCTGGTATCT | CACTTGGTGGTGTCCATCTT |
| Primer set used to amplify *EpRALPH03* also amplified *EpRALPH17*. | | | |
| PCR amplification product was not detected for *EpRALPH08* and *EpRALPH012*. | | | |
| *EpTUB2* and *Ep18S* rRNA primer sequences were obtained from Sharma et al. 2019; *Nb* primer sequences were obtained from Yin et al. 2022 | | | |

| **Table S8**. List of primers used for cloning and Site-directed Mutagenesis (SDM) | | |
|---|---|---|
| Primer Name | Forward Primer (5’-3’) | Reverse Primer (5’-3’) |
| *EpRALPH11R* | AAAAAGCAGGCTTCAACAGGTAC GGATTCAGGTGCGG | AGAAAGCTGGGTCTACGTACTTGC ACTTGGTGTACTCAC |
| *EpRALPH02F* | AAAAAGCAGGCTTCCGCACAACT TTCACTAGCC | AGAAAGCTGGGTCTCATTTTTCATC CGATTCTGT |
| *EpRALPH02R* | AAAAAGCAGGCTTCGCTTACCGC TGTGTTCTTAACG | AGAAAGCTGGGTCTCACTCACAAT TCTTATATTCCTT |
| *FoSRE1R* | AAAAAGCAGGCTTCGCCACTACC TGCGGCAGC | AGAAAGCTGGGTCTTAGCTGCATC CAACAAAGTTGTT |
| *PsFibrillarin2* | AAAAAGCAGGCTTCGCTCCTCCT CCTCGAGGT | AGAAAGCTGGGTCTTACTCAGCAT CTTTCTTCTTCTT |
| *EpRALPH11R$^{H39A}$* | GTTCTGTTTTCCCTGCTGTGGCC ACTGCTTTCAACTTCG | CGAAGTTGAAAGCAGTGGCCACAG CAGGGAAAACAGAAC |
| *EpRALPH11R$^{E52A}$* | CGACGGACCTTACGTCGCTTGGC CTATCACTAGAAAC | GTTTCTAGTGATAGGCCAAGCGAC GTAAGGTCCGTCG |
| *EpRALPH11R$^{R71A}$* | GCAAGTCTAAGAGGGCTATCGTG ATGACCA | TGGTCATCACGATAGCCCTCTTAGA CTTGC |
| *EpRALPH11R$^{R86A}$* | GTTGTTGGAGCTGTGGTGGCACA CAAAGGTGGTGAGTACAC | GTGTACTCACCACCTTTGTGTGCC ACCACAGCTCCAACAAC |

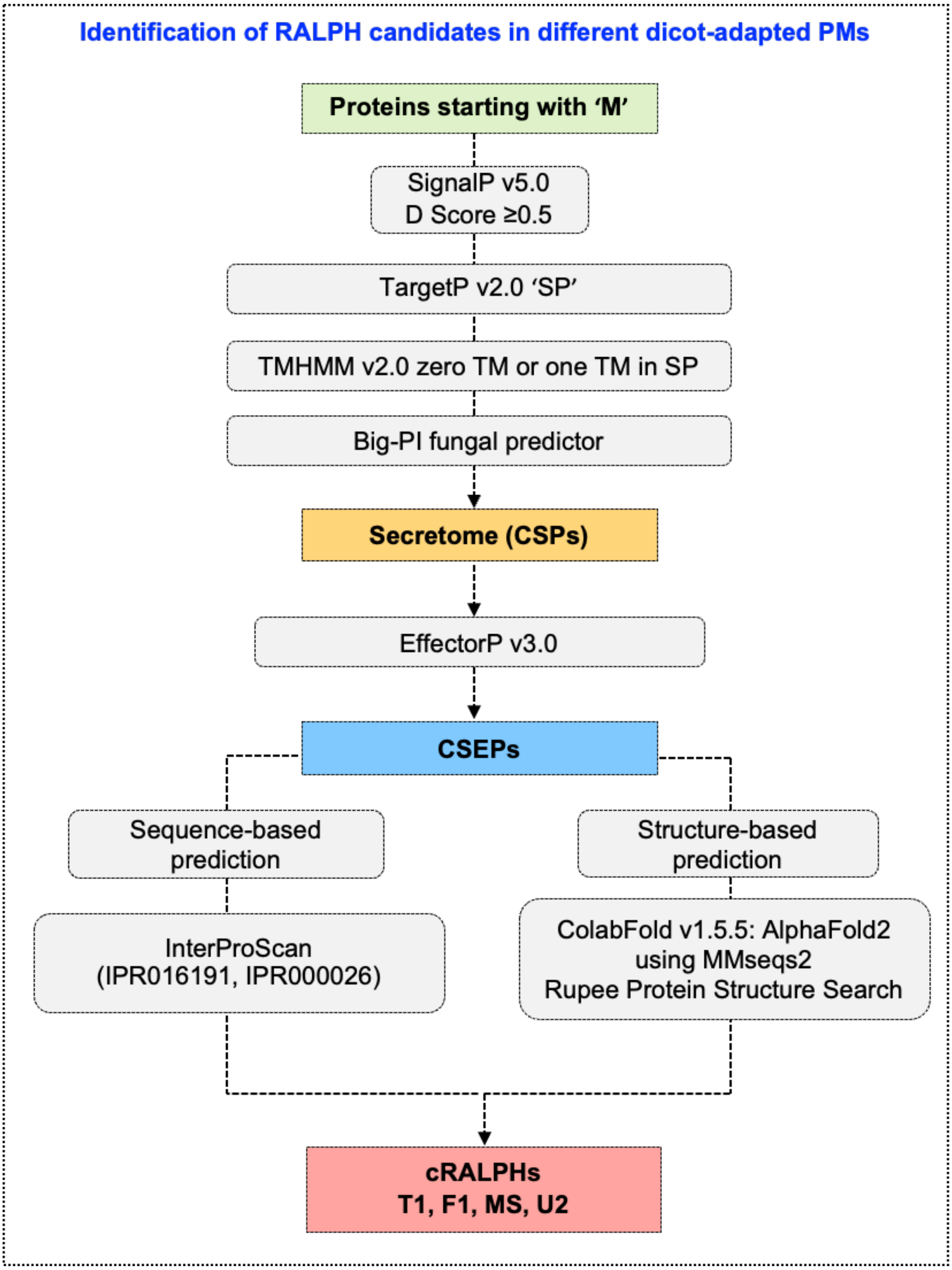


**Figure S1.** Bioinformatics pipeline for predicting RALPH effector candidates in powdery mildew genomes. CSP, candidate secreted proteins; CSEP, candidate secreted effector proteins; cRALPHs, candidate RALPHs

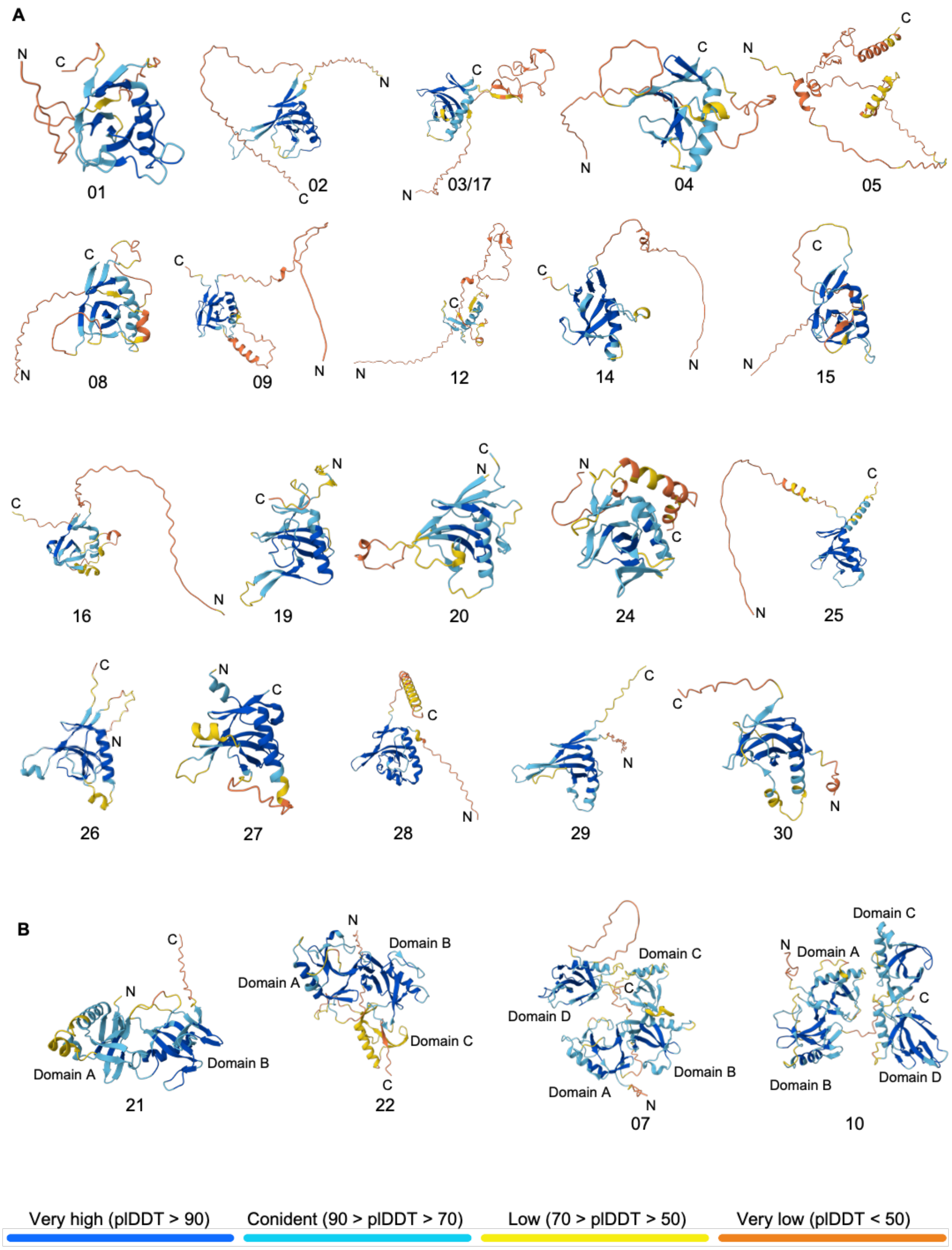


**Figure S2.** AlphaFold 3 (AF3) models of (A) single RNase domain and (B) multi-RNase domain *Ep*RALPH full-length proteins.

**A**

| Coding sequence (CDS) length of multi-RNase domain containing *Ep*RALPHs as presented in the MycoCosm portal | |
|---|---|
| *RALPH ID* | Predicted CDS length (base pairs) |
| *RALPH6* | 1557 |
| *RALPH7* | 1521 |
| *RALPH10* | 1491 |
| *RALPH18* | 738 |
| *RALPH22* | 1038 |
| *RALPH21* | 792 |
| *RALPH23* | 1059 |

**B**

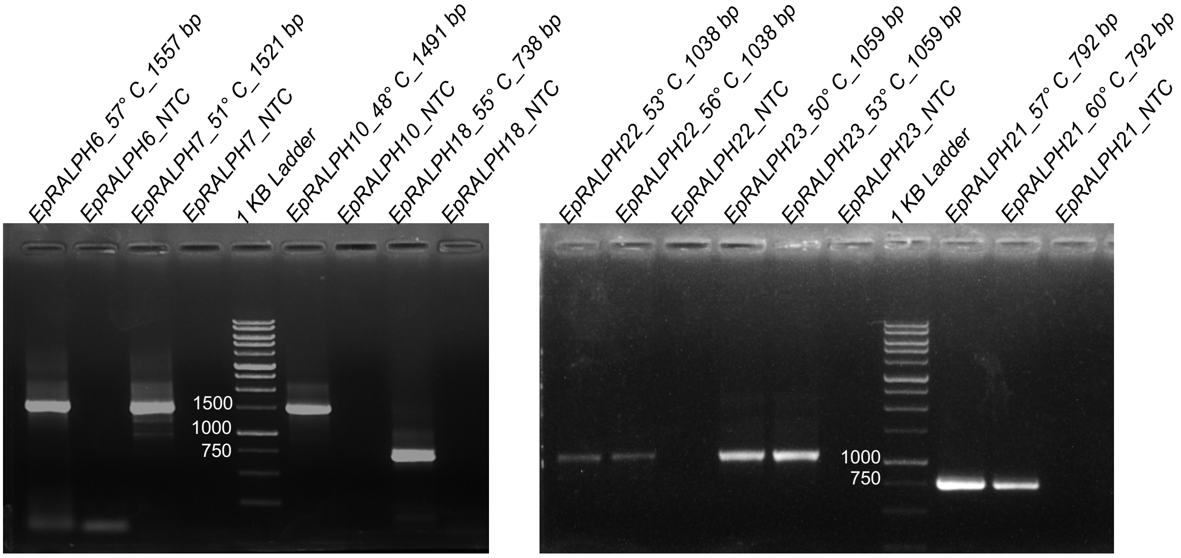


**Figure S3.** Verification of CDS length of multi-RNase domain *EpRALPHs* amplified from *Ep* cDNA versus predicted CDS length provided in MycoCosm. (A) Predicted CDS length of multi-RNase domain *EpRALPH*s (B) PCR amplified CDS of multi-RNase domain *EpRALPH*s run on a 1% agarose gel. Each well is labeled with the *EpRALPH* ID followed by primer annealing temperature and predicted CDS length.

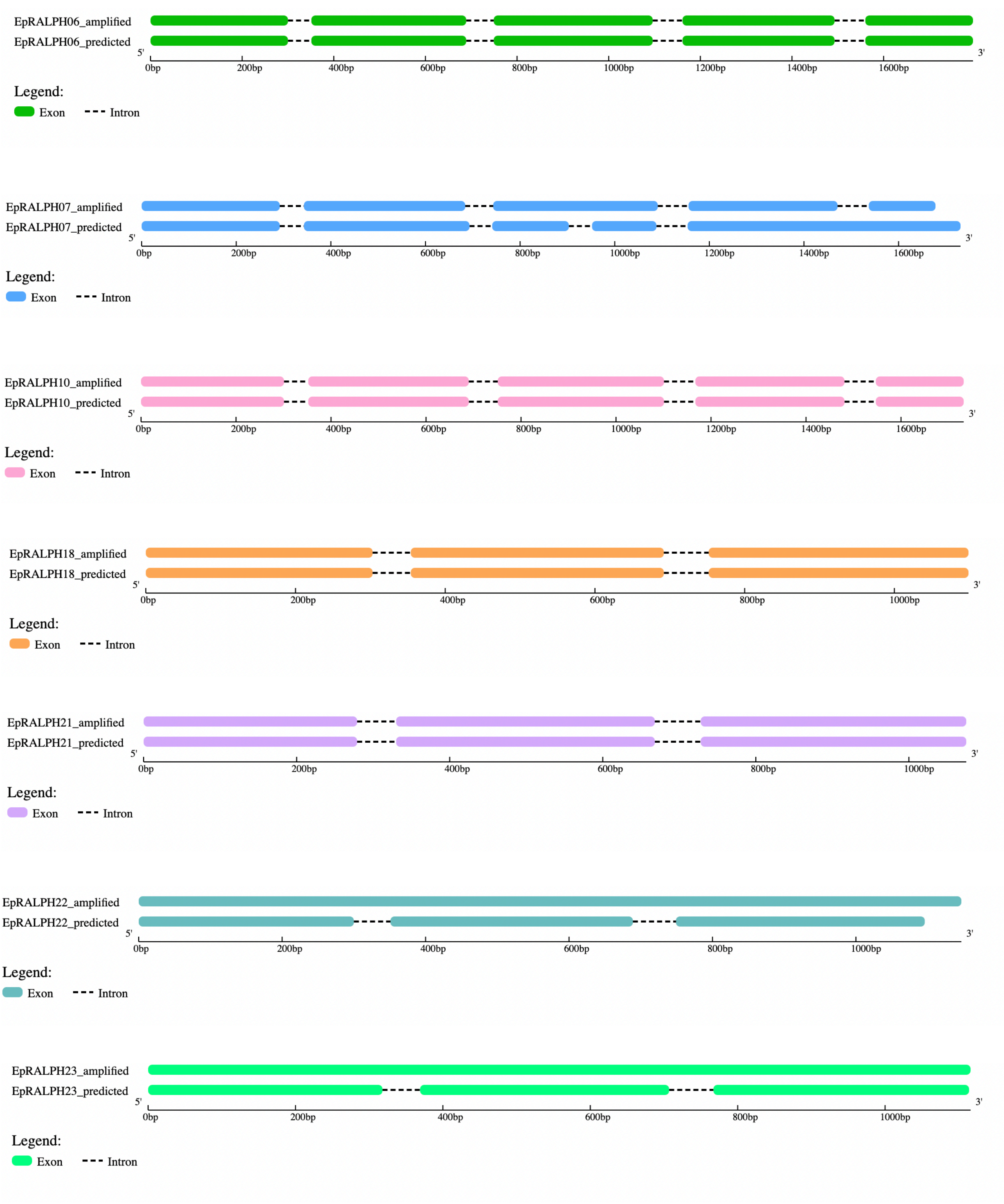


**Figure S4.** The graph represents the number and distribution patterns of exons and introns in the predicted or PCR amplified multi-RNase domain *EpRALPH*s displayed using the Gene Structure Display Server [GSDS2.0, http://gsds.cbi.pku.edu.cn]. Both introns are retained in *EpRALPHs* 22 and 23. The third intron is retained, and a fourth one is introduced in the last exon of *EpRALPH07*.

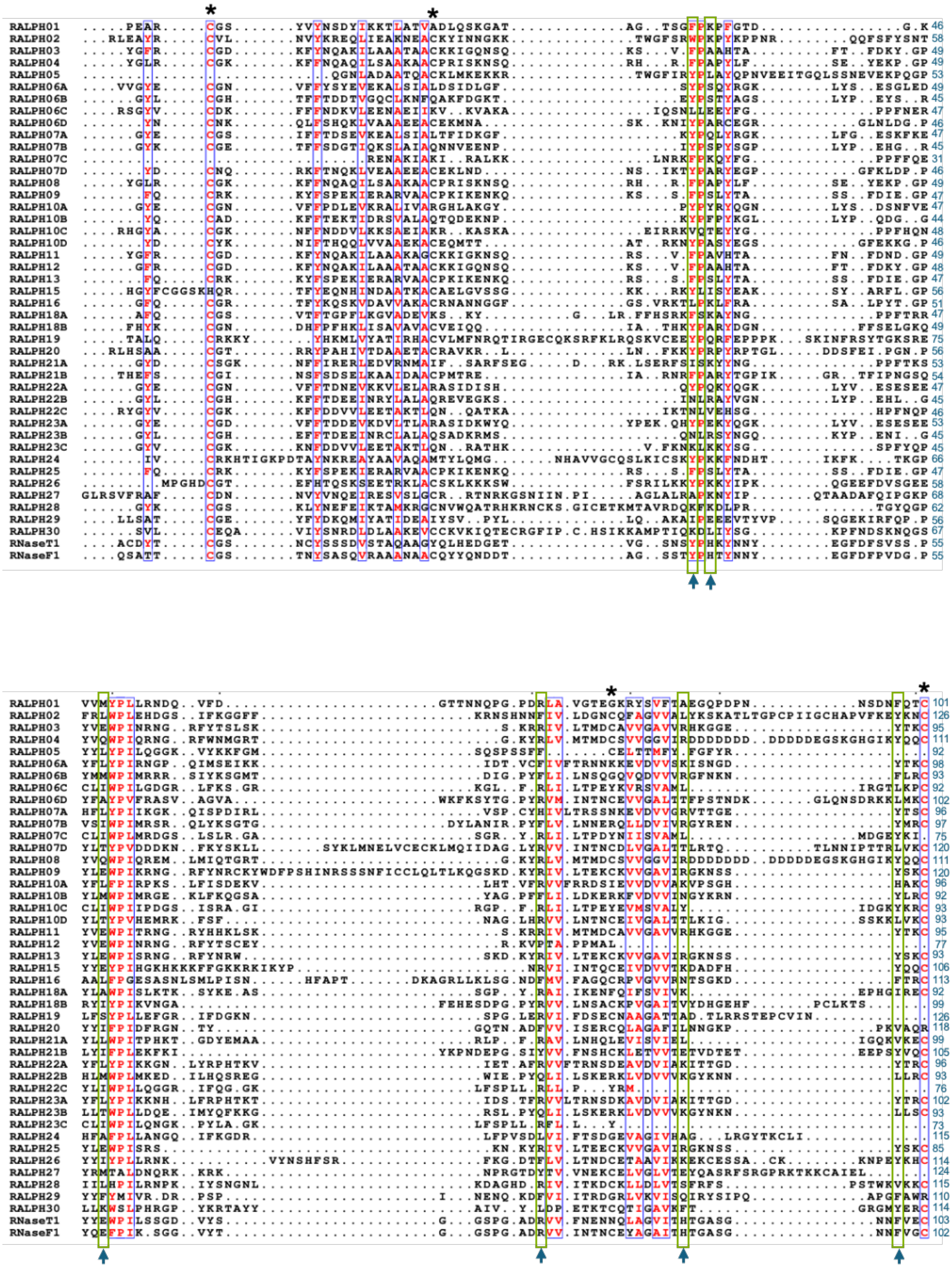

**Figure S5**. Multiple sequence alignment of RNase domains of *Ep*RALPHs along with T1 and F1 RNases, highlighting conservation of cysteine (asterisks) and catalytic residues (arrows). Residues are numbered excluding the signal peptide for all proteins.

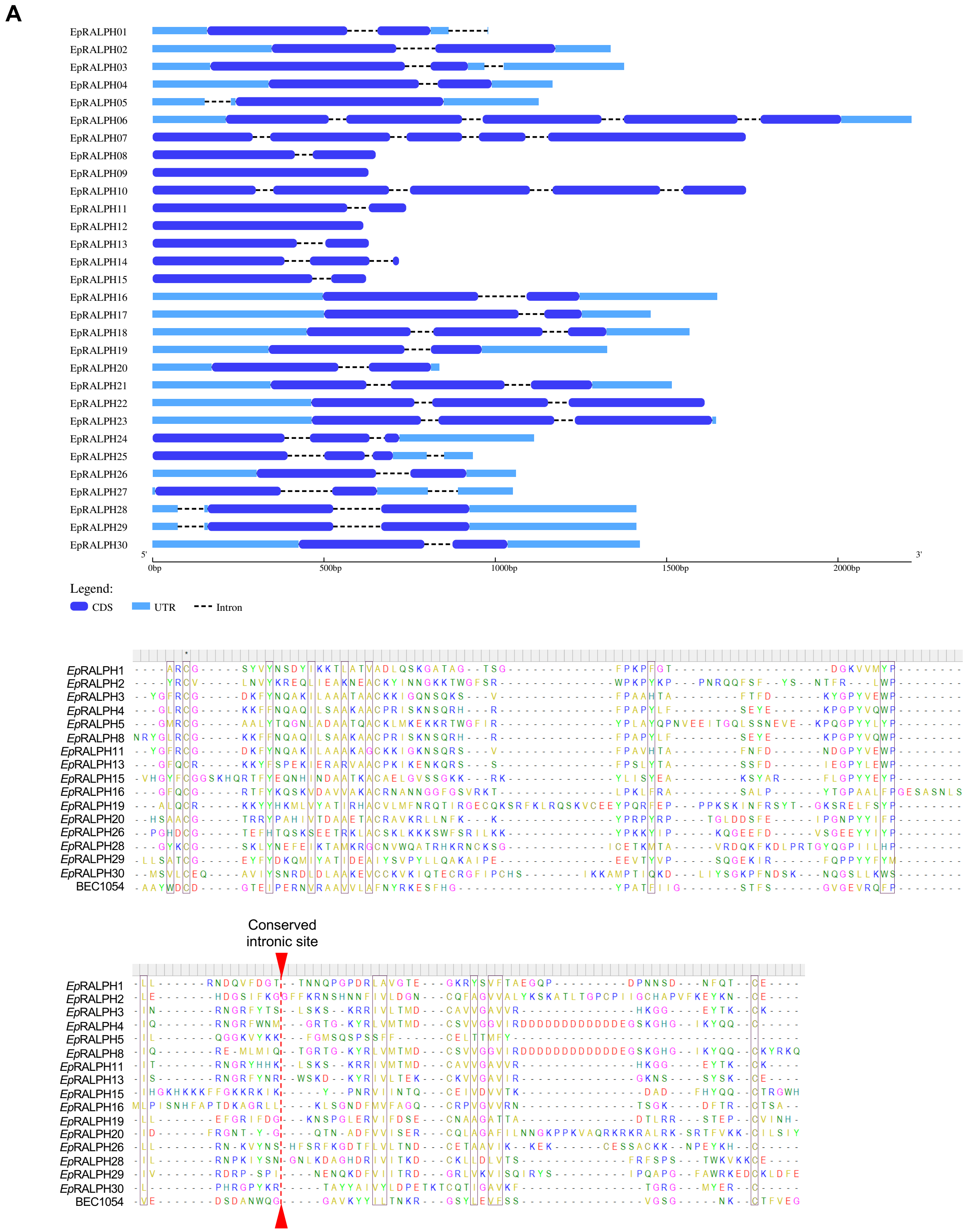


**Figure S6. (A)** The graph represents the number and distribution patterns of exons and introns in the *EpRALPH* genes displayed using the Gene Structure Display Server [GSDS2.0, http://gsds.cbi.pku.edu.cn] **(B)** Multiple sequence alignment of 16 single-intron *Ep*RALPHs and *Bgh*BEC1054 to depict the conserved intronic site (marked with an inverted red triangle), created using the Clustal W algorithm in MEGA12.

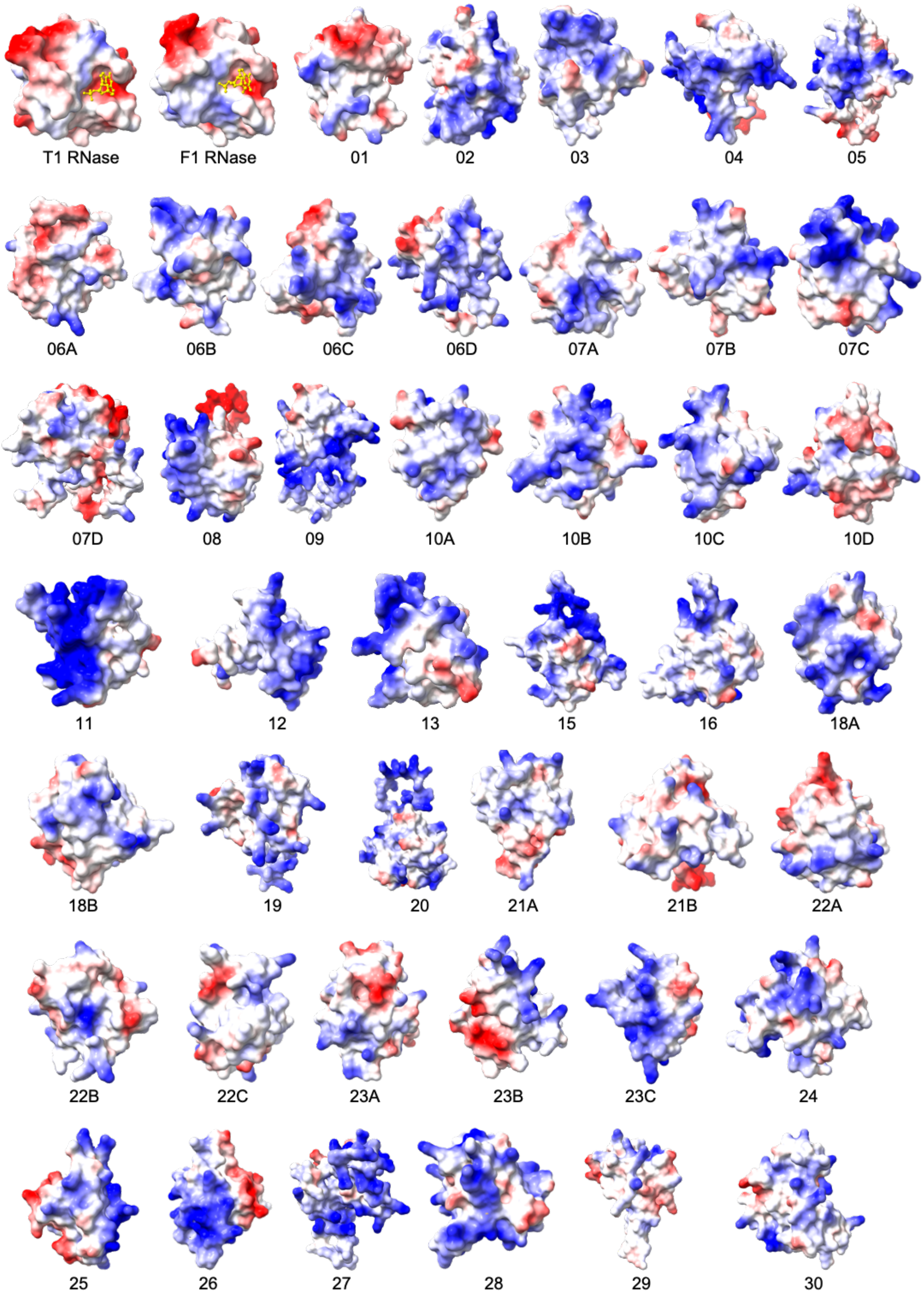


**Figure S7**. Surface charge potential of *Ep*RALPH RNase domains along with T1 (PDB: 1RNT) and F1 (PDB: 1FUT) RNase, generated using ChimeraX. Red, blue, and white indicate acidic (negative), basic (positive), and non-polar or uncharged (neutral) surface residues, respectively, with color intensity corresponding to the strength of basicity and acidity.

**A**

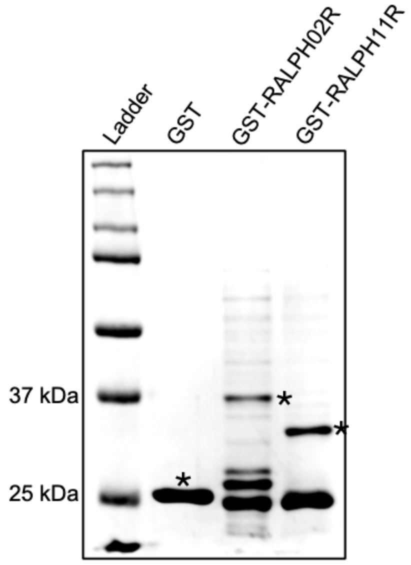


**B**

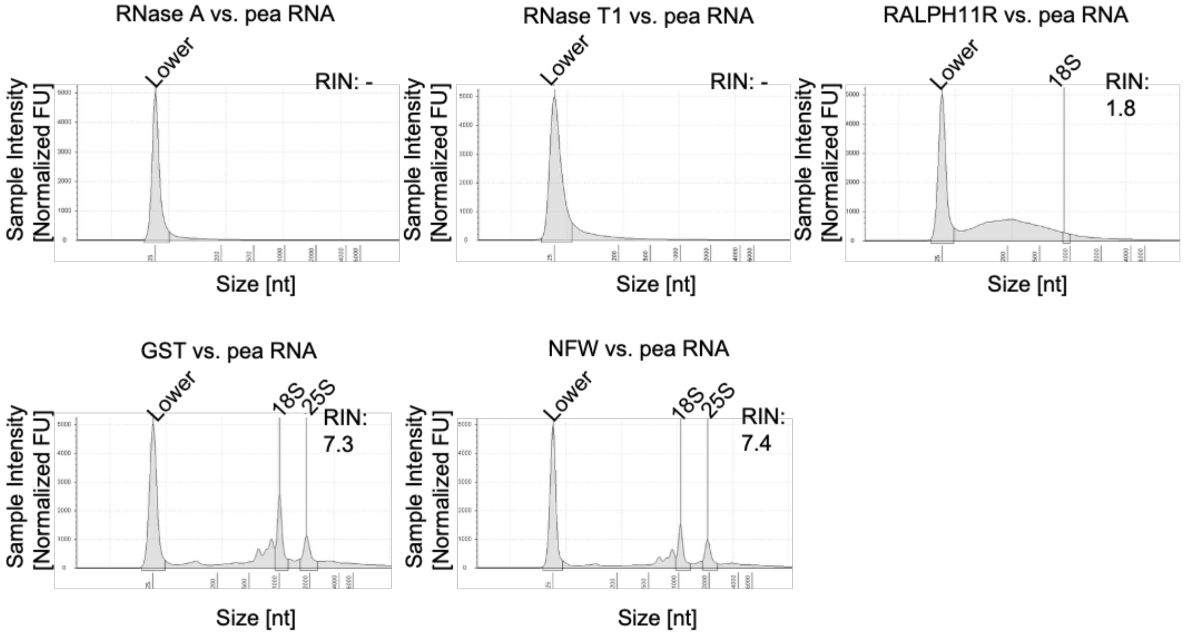


**Figure S8**. (A) SDS-PAGE gel image showing purified GST-RALPH11R, GST-RALPH02R, and GST proteins. One µg of each protein was loaded on a 12% SDS-PAGE gel. (B) TapeStation electropherograms for RNase activity reactions of different proteins with ssRNA (pea RNA) as substrate, included as controls for the *in vitro* RNase assay shown in Figure 7B. RIN, RNA Integrity Number. A higher RIN value indicates better RNA quality and integrity.

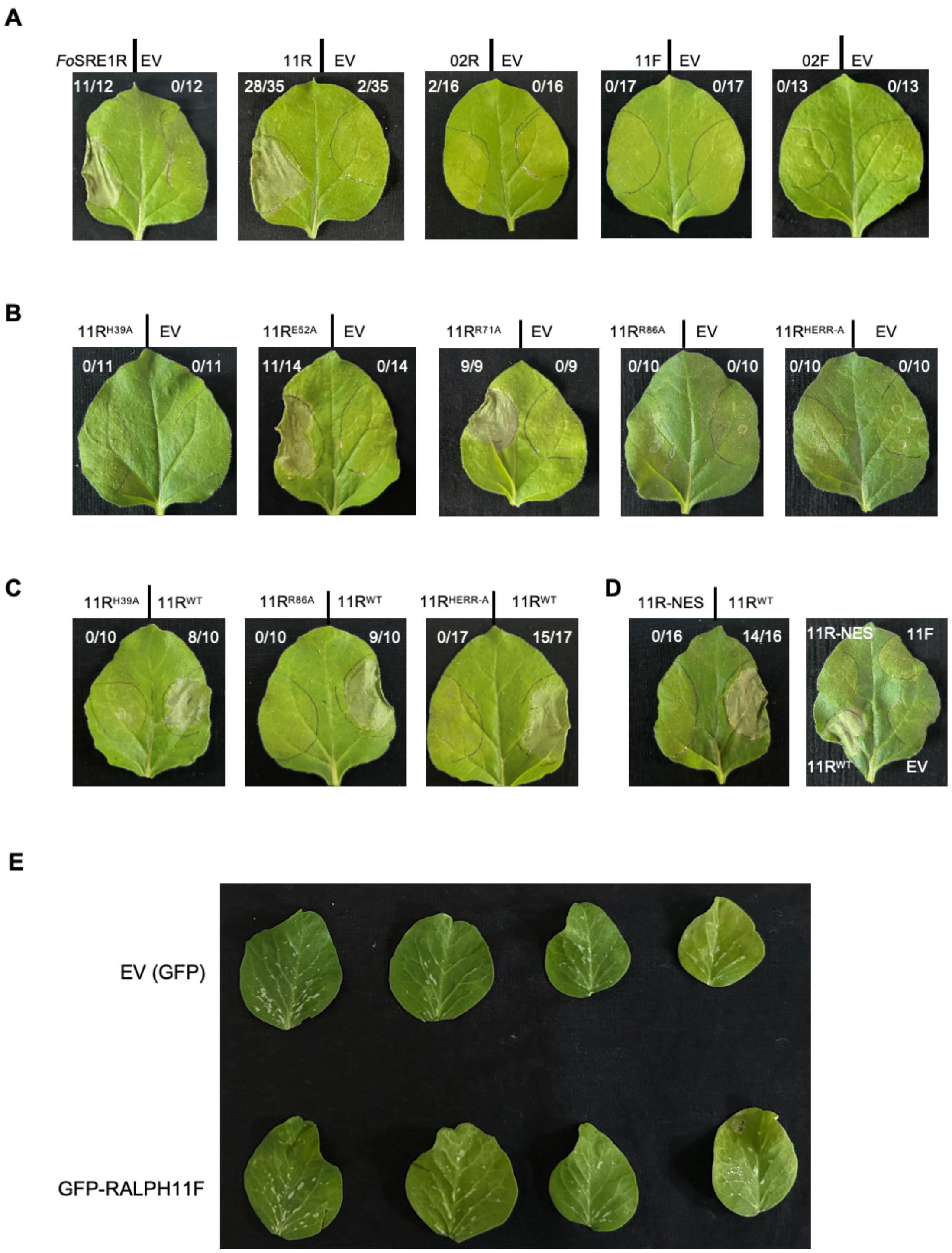


**Figure S9**. Whole leaf images showing presence or absence of cell death symptoms after agroinfiltration of wildtype or mutant constructs of *Ep*RALPHs in (A-D) *Nicotiana benthamiana* and (E) *Pisum sativum* leaves. The number of *N. benthamiana* leaves showing cell death symptoms out of the total number of infiltrated leaves is indicated for each construct. EV, empty vector (negative control); *Fo*SRE1, *F. oxysporum* RNase effector (positive control); 11R and 02R, GFP-*Ep*RALPH11 and 02 RNase domains; 11F and 02F, GFP-*Ep*RALPH11 and 02 full-length proteins minus SP; WT, wildtype; NES, nuclear export signal.
